# The UBP5 histone H2A deubiquitinase counteracts PRC2-mediated repression to regulate Arabidopsis development and stress responses

**DOI:** 10.1101/2022.11.15.516593

**Authors:** James Godwin, Eduardo March, Mohan Govindasamy, Clara Bourbousse, Léa Wolff, Antoine Fort, Michal Krzyszton, Jesús López, Szymon Swiezewski, Fredy Barneche, Daniel Schubert, Sara Farrona

**Author notes:** equal contribution to this work (Eduardo March, Mohan Govindasamy).

## Abstract

Polycomb Repressive Complexes (PRCs) control gene expression through the incorporation of H2Aub and H3K27me3. However, there is limited knowledge about PRCs’ interacting proteins and their interplay with PRCs in epigenome reshaping, which is fundamental to understand gene regulatory mechanisms. Here, we identified UBIQUITIN SPECIFIC PROTEASE 5 (UBP5) as a novel interactor of the PRC2 subunit SWINGER and its associated factor PWO1 in *Arabidopsis thaliana*. As inferred from the functional analyses of *ubp5* CRISPR-Cas9 mutant plants, UBP5 regulates plant development and stress responses, notably by promoting H2A monoubiquitination erasure, leading to transcriptional de-repression. Preferential association of UBP5 at PRC2 recruiting motifs and local H3K27me3 gaining in *ubp5* mutant plants further suggest the existence of functional interplays between UBP5 and PRC2 in regulating epigenome dynamics. In summary, UBP5 provides novel insights to disentangle the complex PRC2 interaction network and is a crucial regulator of the pivotal epigenetic repressive marks H2Aub and H3K27me3.

## Introduction

Histones that form the nucleosome, i.e. basic units of the chromatin, are marked by an array of covalent marks, especially on histone amino terminal tails but also on globular domains. Histone marks impact chromatin structure, modify its packaging and act as an anchor for chromatin-related proteins, transcription factors and other components of the transcriptional machinery ^1^. Therefore, different systems evolved in the eukaryotic nuclei to act as ‘writers’, able to deposit covalent chemical groups on specific histone residues, ‘readers’, which can directly bind and help to interpret histone marks, and ‘erasers’, actively removing histone post-translational modifications. The orchestration of histone modifying enzymes allows for a highly dynamic chromatin regulation crucial to control nuclear structure and transcription ^2^. Two important histone modifications that are well conserved between plants and animals are the trimethylation on the lysine 27 of the histone H3 (H3K27me3) ^3^ and the monoubiquitination of the histone H2A that in plants mostly occurs on the lysine 121 (H2Aub) ^4^.

H3K27me3 and H2Aub are deposited, both in plants and animals, by two major types of Polycomb repressive complexes (PRCs), respectively PRC2 and PRC1. PRC2 is a four-core subunit complex in which the catalytic component is a SET (Su(var), Enhancer of zeste, Trithorax) domain histone methyltransferase (HMT) ^5, 6^. Analyses in different plant genomes showed that PRC2 decorates approximately 20-25% of euchromatic genes with H3K27me3, which switches them off in response to internal and external cues ^7, 8^. In plants, PRC1 is formed by E3 ligases and other auxiliary proteins ^5, 9^. Both PRCs maintain an intricate relationship in which members of the two complexes can directly interact, have common associated proteins and share target genes. This is also reflected in their activities as H3K27me3 can precede H2Aub (i.e. hierarchical model) or oppositely follows this modification on the chromatin. Furthermore, both marks can independently regulate different set of genes ^7, 9^.

In animals, H2AK119ub can be erased by the Polycomb Repressive-Deubiquitinase (PR-DUB) complex ^10^. This complex contains a DUB protein of the ubiquitin carboxy-terminal (UCH) family, which does not have an obvious orthologous in plants ^11^. Indeed, the PR-DUB has not been described in plants so far, but two proteins of the UBIQUITIN PROTEASE (UBP) family, UBP12 and UBP13 redundantly mediate H2A deubiquitination ^12, 13^ and interact with LIKE HETEROCHROMATIN PROTEIN 1 (LHP1) ^12^, a H3K27me3 reader and interactor of both PRC2 and PRC1 components ^7, 9^. UBP12/13 regulate a similar set of genes with PRC2 and PRC1 ^13^.

To develop their activities, PRCs require a complex network of protein-protein interactions ^7^. We and others recently demonstrated that PWWP-DOMAIN INTERACTOR OF POLYCOMBS1 (PWO1) is a key regulator of PRC2 activity, able to interact with the HMTs of the PRC2 complex ^14^ and to form part of the PEAT complex (<u>P</u>WO/PWWP-<u>E</u>PCRs (ENHANCER OF POLYCOMB RELATED)-<u>A</u>RIDs (AT-RICH INTERACTION DOMAIN-CONTAINING)-<u>T</u>RBs (TELOMERIC REPEAT BINDING)) involved in heterochromatin dynamics ^15^. Still, we are far from understanding the molecular impact of the PWO1-PRC2 interaction.

Here we show that UBP5 is a novel interactor of PRC2 and PWO1 that is able to affect both H3K27me3 and H2Aub marks as well as the expression of a set of PRC2 target genes in *Arabidopsis thaliana* (Arabidopsis). Telobox and GAGA motifs, previously related to PRC2 recruitment ^16, 17^, are among the most enriched signatures of UBP5 binding to the chromatin. The vast majority of UBP5 direct target genes showed either hyper-marking or *de-novo* marking by H2Aub in *ubp5* plants, altogether indicating that UBP5 acts as a sequence-specific eraser of this epigenetic mark. Together, our data uncovers UBP5 as a new PRC2-interactor module directly controlling H2Aub deubiquitination and affecting H3K27 trimethylation to regulate gene expression.

## Results

### UBP5 is a novel interactor of PRC2 and PWO1

We had identified the UBIQUITIN PROTEASE 5 (UBP5) protein as the most abundant interactor co-immunoprecipitated with Arabidopsis PWWP-DOMAIN INTERACTOR OF POLYCOMBS1 (PWO1) ^18^. Furthermore, data mining of proteins in co-immunoprecipitation (co-IP) experiments with PEAT components also identified UBP5 ^15^. Therefore, we aimed to understand the link between UBP5, PWO1 and PRC2. Firstly, to elucidate the sub-cellular localisation of UBP5, transient inducible expression was performed using the β-estradiol– inducible 35S promoter (*i35S*) fused to an *UBP5* (*i35S::UBP5-GFP*) construct in *Nicotiana benthamiana* (*N. benthamiana*) and found that UBP5 is exclusively nuclear, localises all over the nucleoplasm in a diffused way but not in the nucleolus (Fig. 1A). Further, we analysed the possibility of an interaction between UBP5 and PWO1 *in planta*. Using a similar approach, we co-expressed PWO1-GFP and UBP5-mCherry fusion proteins in *N. benthamiana*. It is noteworthy that, as previously shown for CLF, co-expression of both proteins modified UBP5 localisation recruiting it to PWO1-containing nuclear speckles and, to a lower extent, co-localisation of both proteins was also observed all over the nucleoplasm (Fig. 1B) ^14, 18^. PWO1-UBP5 association in both speckles and nucleoplasm was demonstrated by Foster resonance energy transfer with acceptor photobleaching (FRET-APB). FRET-APB efficiencies for co-expressed samples were significantly higher than the negative controls (PWO1-GFP and UBP5-GFP expressed without donor mCherry construct) (Fig. 1C). The FRET-APB donor signal intensity was significantly higher in the speckles than in the nucleoplasm, which can be due to PWO1 and UBP5 stronger association and/or because of a higher probability of contacts between both proteins within the speckles (Fig. 1C). Yeast two-hybrid (Y2H) assays not only confirmed the interaction of UBP5 with PWO1 but also revealed its interaction with the PRC2 HMT subunit SWINGER (SWN) ΔSET (SWN clone lacking the SET domain;^19^) (Fig. 1D). *In planta* interaction between SWNΔSET and UBP5 was further confirmed using co-IP assays in *N. benthamiana* (Fig. 1E). Therefore, UBP5 is an interactor of PWO1-PRC2 suggesting the possibility that it may play a role in PRC-mediated regulation of gene expression. Furthermore, Y2H assays showed interaction of UBP5 with EMBRYONIC FLOWER 2 (EMF2), another PRC2 component ^20^, which further confirms the PRC2-UBP5 connection (Supplementary Fig. 1).

**Figure 1.**
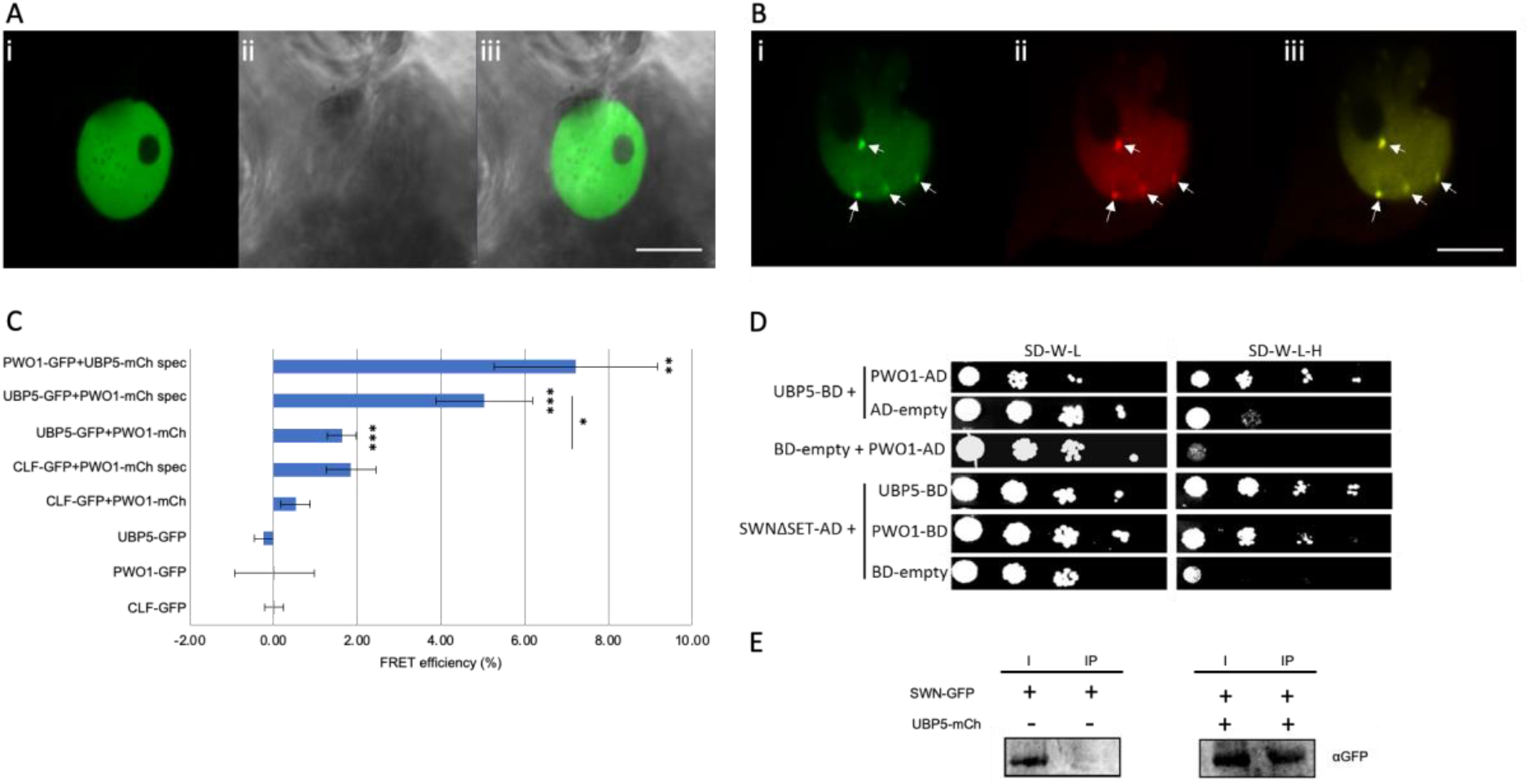
UBP5 is a nuclear protein that interacts with PRC2 and colocalises with PWO1. A and B, transient and inducible expression in *N. benthamiana* epidermal cells, bar = 10µm. A, *i35S::UBP5-GFP* (i, confocal; ii, bright field; iii, overlay). B, *i35S::UBP5-GFP* and *i35S::PWO1-mCherry* co-transformation (i, *i35S::UBP5-GFP;* ii, *i35S::PWO1-mCherry;* iii, overlay). Arrows indicate speckles. C, FET-APB measurements for nuclei exemplified in B, with a distinction for speckle (spec) and non-speckle localisation. CLF-GFP and PWO1-mCherry measurement was used as positive control (Mikulski et al., 2019). An average of efficiency for n = 7-19 is shown. Significance level was measured in comparison to control or as indicated using Student’s t-test and is represented by *p<0.05, **p<0.01, ***p<0.001. D, Y2H analyses confirm UBP5-PWO1 interaction and show an UBP5-SWN interaction. Yeast cells containing the different construct combinations on selective medium for plasmids (-LW; - leucine, tryptophan) or for reporter gene activation (-LWAH; -leucine, tryptophan, adenine, histidine). Serial solutions were used. BD, GAL4-DNA binding fusion; AD, GAL4-DNA activation domain fusion. SWNΔSET, SWN construct lacking the SET domain. E, Co-IP analyses confirming SWN-UBP5 interaction. IP was performed with anti-mCherry antibody and proteins were detected by western blot with anti-GFP. I, 5% input; IP, immunoprecipitation.

### UBP5 is an essential plant developmental and stress responses regulator

To understand UBP5 molecular functions in Arabidopsis, we generated an *ubp5* deletion mutant line via the CRISPR/Cas9 system with two guide RNAs, which partially deleted both DUSP and UBP conserved domains (Supplementary Fig. 2A-C). The phenotypic analyses of *ubp5* mutant plants showed pleiotropic defects such as stunted growth due to the lack of apical dominance (Fig. 2A (i-iii)), shorter roots and hypocotyl length (Fig. 2A ii and 2B), floral architecture defects (Fig. 2A (v-vi)), fertilisation defects (Supplementary Fig. 2D) and poor pollen germination (Supplementary Fig. 2E), suggesting that UBP5 acts as a developmental regulator at different stages of the plant life cycle. Stable transformation of *UBP5pro::UBP5-eGFP* was able to fully rescue the developmental pleotropic phenotypes of *ubp5* (Fig. 2A (iv)). qRT-PCR analyses further showed no significant difference in the relative expression of *UBP5* between Col-0 and the complementation line *UBP5pro::UBP5-eGFP;ubp5* (Supplementary Fig. 3A-B). Transcriptional analyses of *ubp5* seedlings showed that 345 genes were up-regulated, and 478 genes were down-regulated (Fig. 2C; Supplementary list 1). Mis-regulation of major developmental genes including *KNOTTED-LIKE FROM ARABIDOPSIS THALIANA* (*KNAT1*), *PISTILLATA, MERISTEM DISORGANIZATION 1* (*MDO1*), *SAMBA* and *GAMETOPHYTIC DEFECTIVE 1* (*GAF1*) correlated with some of the observed *ubp5* mutant phenotypes (Supplementary list 2). In addition, considering the bushy-like phenotype, we analysed the expression of several genes encoding transcription factors involved in controlling the shoot apical meristem that are also PRC2 repressed (i.e., marked by H3K27me3). Our RT-qPCR analyses demonstrated their upregulation in *ubp5* (Supplementary Figure 4A-E). Gene Ontology (GO) analyses identified that genes associated with biotic and abiotic stress responses terms were significantly enriched among all *ubp5* mis-regulated genes (Fig. 2D). Consistently with previous studies showing that PRC2-associated components do not only regulate expression of genes related to plant development ^13, 21, 22^, our results indicate a dual role of UBP5 in regulating both Arabidopsis developmental and stress responses.

**Figure 2.**
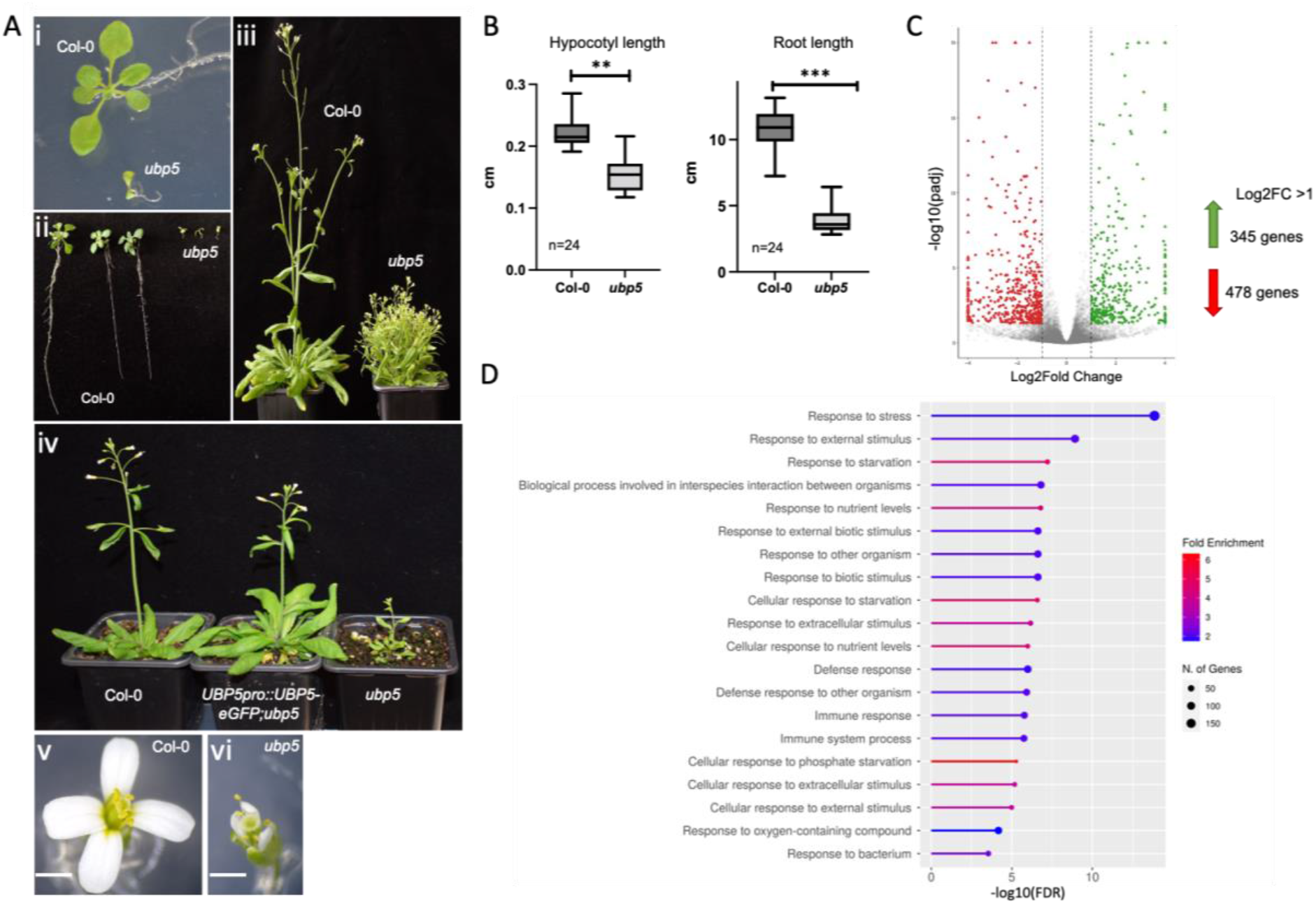
UBP5 is an essential plant developmental and stress regulator. A, Phenotypic characterisation of *ubp5* mutant line: i, smaller seedlings; ii, shorter primary roots; and iii) stunted and bushy growth (Note: in iii *ubp5* plant was 2 weeks older than the *Col-0* plant) compared to Col-0 plants; iv, complementation of *ubp5* mutant phenotypes in 4-week-old Arabidopsis plants (an *UBP5pro*::*UBP5-eGFP* construct was used for the complementation of *ubp5*); v, floral phenotype of Col-0 and vi, *ubp5*. Bar *=* 1 mm. B, Hypocotyl and root length of *ubp5* versus Col-0 measured after 10 days post-germination. Error bars represent standard deviation, significance tested using student t-test, **p < 0.05, *\*\*\**p< 0.001. C, MA Scatter plot of log2FC versus the log10 basemean. Genes with a p adjusted value (padj) lower than 0.05 are colored. The genes with Log2FC <1 or Log2FC < –1 (padj < 0.05) were considered for further analysis. D, Functional categorisation of *ubp5* mis-regulated (upregulated and downregulated) genes based in ShinyGO v0.75 analysis. GO analysis of *ubp5* mis-regulated genes based on biological process with False Discovery Rate (FDR) < 0.05.

**Figure 3.**
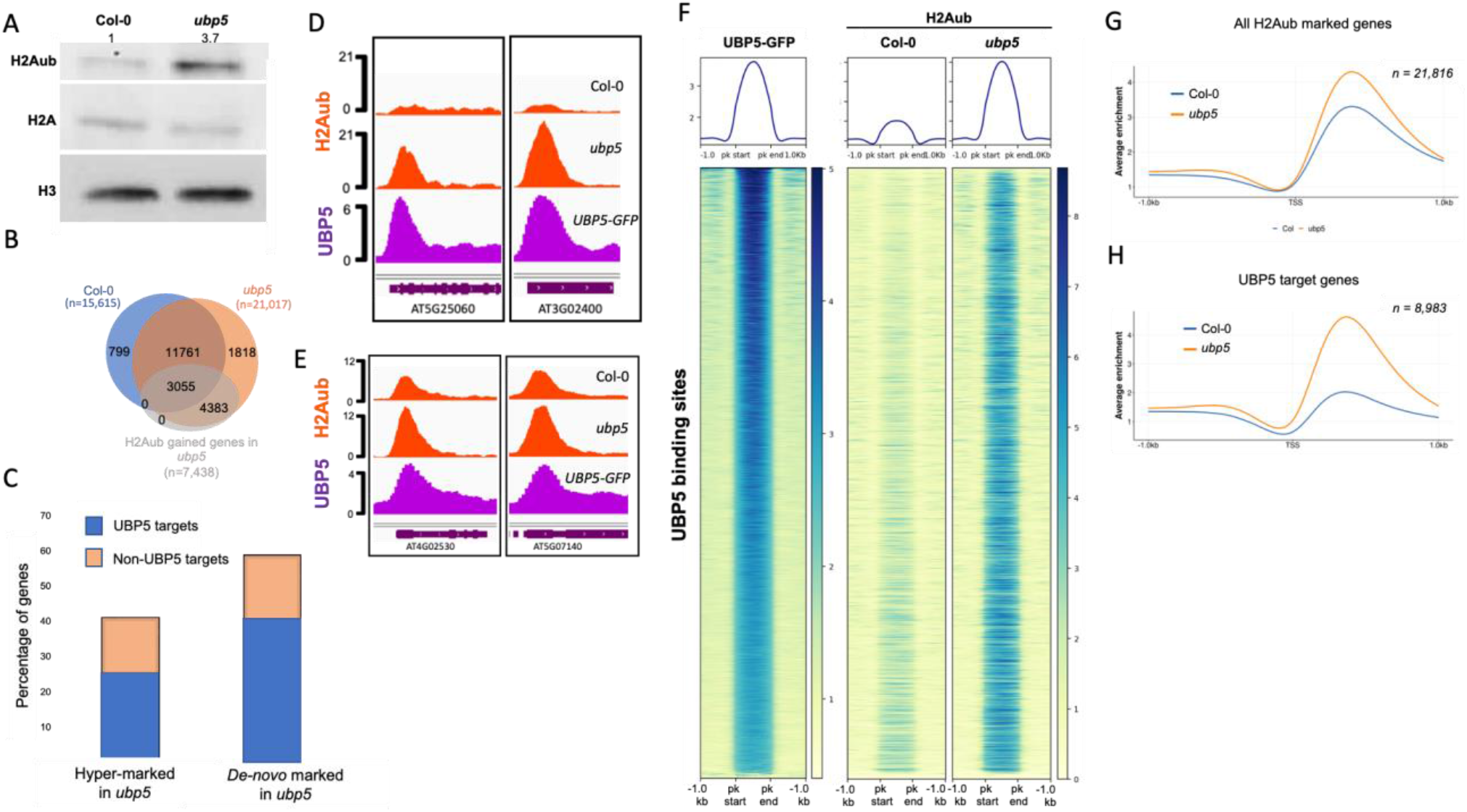
UBP5 acts as a H2A deubiquitinase. A, Western blot of H2Aub and H2A levels in histone extracts from seedlings of Col-0 and *ubp5*. Histone H3 is used as loading control. The numbers above the gel lanes represent the relative H2Aub levels, which was determined from the band intensity using ImageJ software. B, Venn diagram showing the overlap between H2Aub marked genes in Col-0, *ubp5* and H2Aub gained genes in *ubp5*, n represents the number of genes. The genes are considered as marked when an overlapping H2Aub peak is present in at least two biological replicates based on MACS3 peak calling (q <0.05 and score >30) and H2Aub gained genes in *ubp5* were found using DESeq2 analysis (FDR < 0.05). C, The graph represents the two categories of genes showing H2Aub changes in *ubp5:* hyper-marked genes –genes that show a hyper enrichment of H2Aub in *ubp5* if they were already marked in the Col-0– and d*e-novo* marked genes –genes only marked in the *ubp5* but not in the Col-0. Differential H2Aub analysis was done using DESeq2 analysis (FDR < 0.05). D-E, IGV browser views of representative UBP5 target loci where (D) *de-novo* marked genes in the *ubp5* mutant and (E) H2Aub is hyper-marked in *ubp5*. Gene structures and names are shown underneath each panel. F, Heatmaps showing H2Aub distribution on genomic sequences targeted by UBP5 for Col-0 and *ubp5*. UBP5 binding peaks are clustered based on higher to lower enrichment from top to bottom. G, Metagene plot of average H2Aub distribution over 1 kb upstream and downstream from the transcription start site (TSS) of all the H2Aub marked genes in Col-0 and *ubp5*. H, Metagene plot of average H2Aub distribution over UBP5 target genes in Col-0 and *ubp5*.

**Figure 4.**
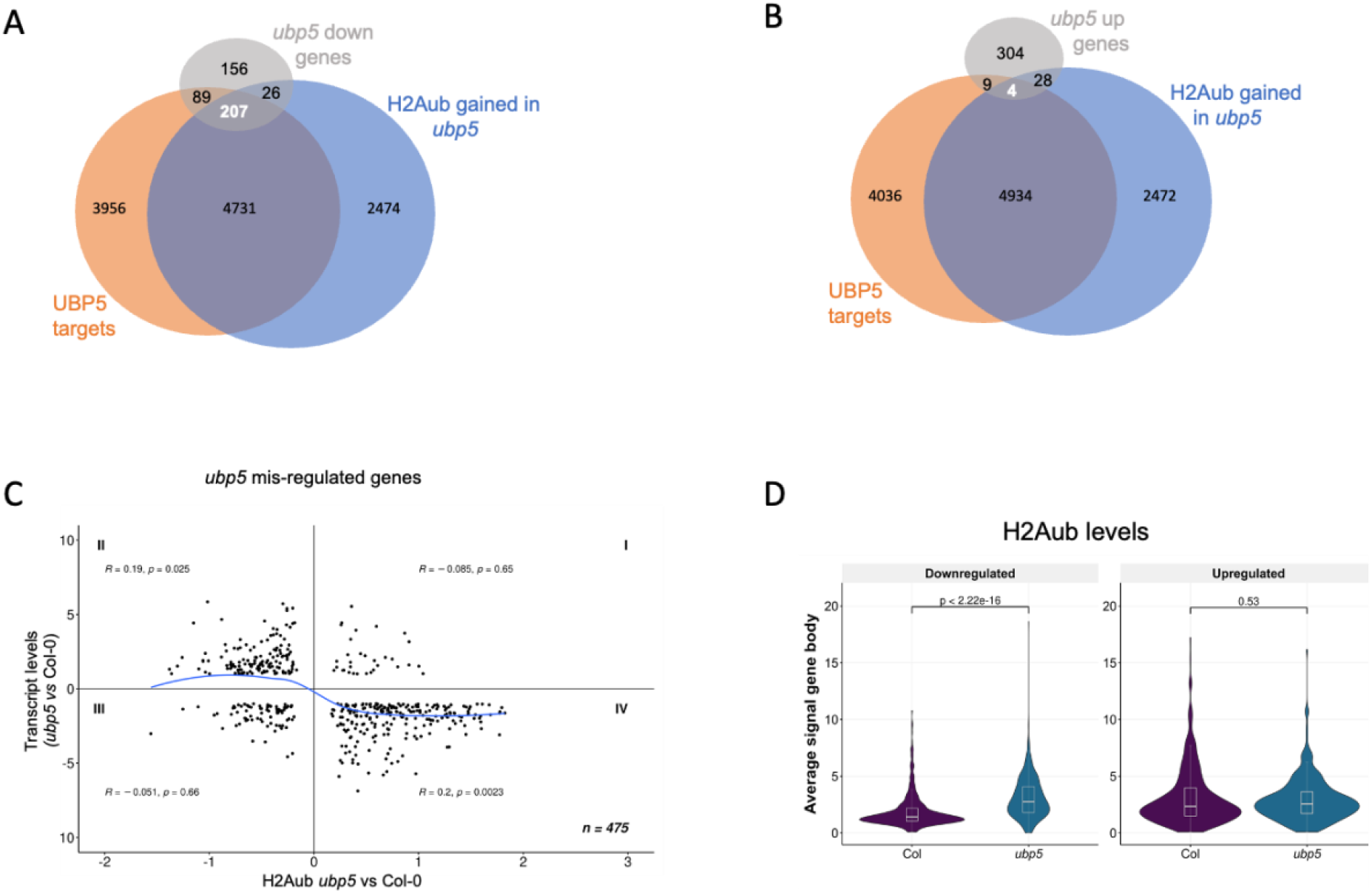
UBP5 mediates transcriptional de-repression. A-B, Venn diagrams showing UBP5 targets, H2Aub gained genes in *ubp5* and (A) downregulated or (B) upregulated genes in *ubp5* mutant. C, Scatter plot showing the correlation between H2Aub and gene expression changes between Col-0 and *ubp5* plants. The x-axis shows Log2FC levels of H2Aub marked genes as determined by DESeq2 analysis (FDR < 0.05). The y-axis shows expression Log2FC of misregulated genes in *ubp5* as determined by DESeq2 (>1 fold variation, FDR < 0.05). For each quadrant, the correlation coefficient (R) along with the significance (p values) are shown. The blue curve shows trend-line from LOWESS smoother function. Quadrant IV shows higher correlation between low expressed genes and hyper-marking of H2Aub. D, Violin cum box plots represents the average signal of H2Aub at gene body for downregulated and upregulated genes in Col-0 and *ubp5*. The median (middle line), upper and lower quartiles (boxes) are indicated. Statistical significance is tested according to one-sided Mann–Whitney– Wilcoxon test, p values are indicated above the plot.

### UBP5 deubiquitinates H2A

UBP5 was shown *in vivo* to be involved in de-ubiquitination of hexa-ubiquitin substrates ^23^ and other UBP family members have been linked to the histone monoubiquitination removal ^24, 25, 26^. In addition, the existence of the interaction between UBP5, PRC2 HMTs and PWO1 made us speculate that UBP5 may contribute to PRC-mediated histone monoubiquitination dynamics. Therefore, we analysed different histone marks abundance in *ubp5* and Col-0 seedlings by western blot (WB) assays and, in good agreement with UBP5 acting in H2Aub removal, we found that H2Aub bulk levels were more than 3-fold higher in *ubp5* (Fig. 3A). To gain insight into the affected loci, we profiled the genome-wide distribution of H2Aub in *ubp5* and Col-0 seedlings using ChIP-seq. Our H2Aub data in Col-0 seedlings showed a good overlap with previous published data (Supplementary Fig. 5) and, when compared to Col-0 seedlings, we observed a large increase in the number of genes uniquely marked by H2Aub in *ubp5* (21,017 in *ubp5* instead of 15,615 genes in Col-0; Supplementary list 3-4), which includes genes that differentially gained H2Aub in *ubp5* (n=7,438; Fig. 3B), hence UBP5 is necessary to erase or decrease H2Aub in several thousands of genes.

**Figure 5.**
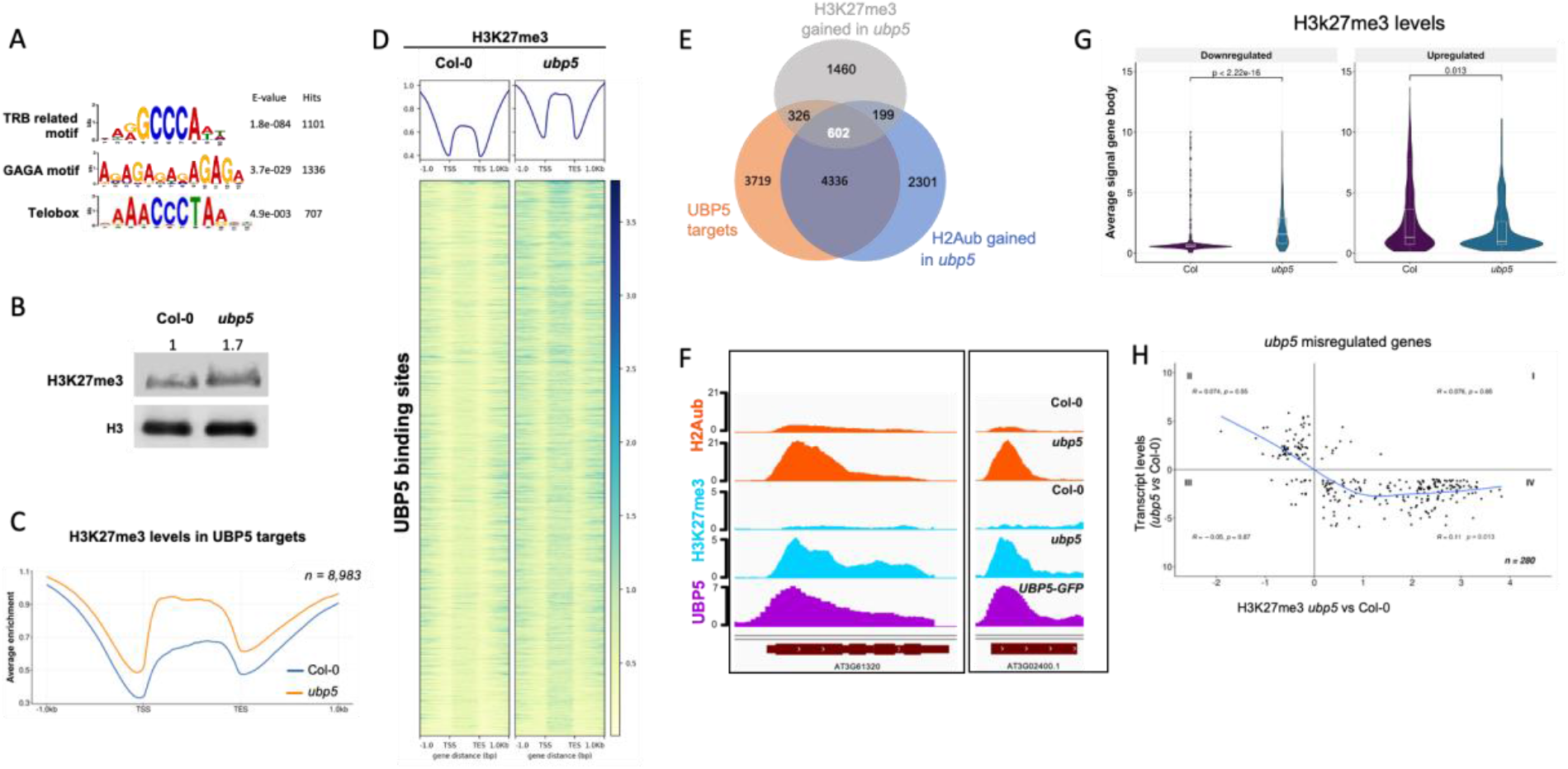
UBP5-mediated H2Aub deubiquitination prevents deposition of H3K27me3. A, Motif enrichment analysis of UBP5 target genes. The sequence logos, accuracies and hits of the best motifs found by MEME-ChIP (Bailey, 2021). B, Western blot of H3K27me3 levels in histone extracts from seedlings of Col-0 and *ubp5*. Histone H3 is used as loading control. The numbers above the gel lanes represent the relative H3K27me3 level, which was determined from the band intensity using ImageJ software. C, Metagene plot of average H3K27me3 enrichment over the UBP5 target genes in Col-0 and *ubp5*. D, Heatmap showing the distribution of H3K27me3 on UBP5 binding sites for Col-0 and *ubp5*. UBP5 binding peaks are clustered based on higher to lower enrichment from top to bottom. E, Venn diagram representing the overlap between UBP5 targets, H2Aub and H3K27me3 gained genes in *ubp5* (padj <0.05) as determined by DESeq2. F, IGV browser snapshots of representative UBP5 target genes in which H2Aub and H3K27me3 are gained in the *ubp5* mutant. Gene structures and names are shown underneath each panel. G, Violin cum box plots represents the average signal of H3K27me3 at gene body for downregulated and upregulated genes in Col-0 and *ubp5*. The median (middle line), upper and lower quartiles (boxes) are indicated. Statistical significance is tested according to one-sided Mann– Whitney–Wilcoxon test, p values are indicated above the plot. H, Scatter plot showing the correspondence between H3K27me3 and gene expression changes in between Col-0 and *ubp5* plants. The x-axis shows Log2FC levels of H3K27me3 marked genes as determined by DESeq2 analysis (FDR < 0.05). The y-axis shows expression Log2FC of mis-regulated genes in *ubp5* as determined by DESeq2 (>1 fold variation, FDR < 0.05). The blue curve shows trend-line from LOWESS smoother function. The correlation coefficient (R) along with the significance (p values) are shown. Quadrant IV shows significant correlation between low expressed genes and gaining H3K27me3.

To test whether UBP5 could act in H2Aub removal in *cis*, we further analysed the genome-wide association of UBP5-GFP in our *UBP5pro::UBP5-eGFP;ubp5* line. Notably, UBP5 binding extends to a large part of the plant genome since the UBP5-GFP ChIP-seq profiling identified 8,983 genes as direct targets of UBP5 (Supplementary Fig. 6A-C; Supplementary list 5), which corresponds to ∼27% of the total number of Arabidopsis genes according to TAIR 10 annotation ^27^. More precisely, UBP5 directly targets 69% of the genes gaining *de novo* a H2Aub peak in *ubp5* (i.e., *de-novo* marked genes, Fig. 3C and 3D), and 61% of the genes for which H2Aub peaks are increased in *ubp5* (i.e., hyper-marked genes) (Fig. 3C and 3E). Importantly, there is a sharp co-localisation between UBP5 chromatin association and domains where the H2Aub mark was gained in *ubp5* (Fig. 3D-E; Supplementary Fig. 7A; Supplementary list 6). This frequent co-occurrence strongly argues in favour for a direct role of UBP5 in H2Aub deubiquitination at its binding sites (Fig. 3F). Further supporting this observation, increase in H2Aub levels in *ubp5* is more evident at UBP5 target genes than for other, non-targets, H2Aub marked genes (Fig. 3G-H and Supplementary Fig. 7B). To confirm these observations, selected UBP5 targets that are H2Aub hyper-marked in *ubp5* were further validated by ChIP–qPCR (Supplementary Fig. 7C). Overall, these results indicate that UBP5 acts in *cis* on H2Aub mark by both maintaining the H2Aub level in a set of genes marked with this modification and erasing the H2Aub mark from a larger set of genes.

**Figure 6.**
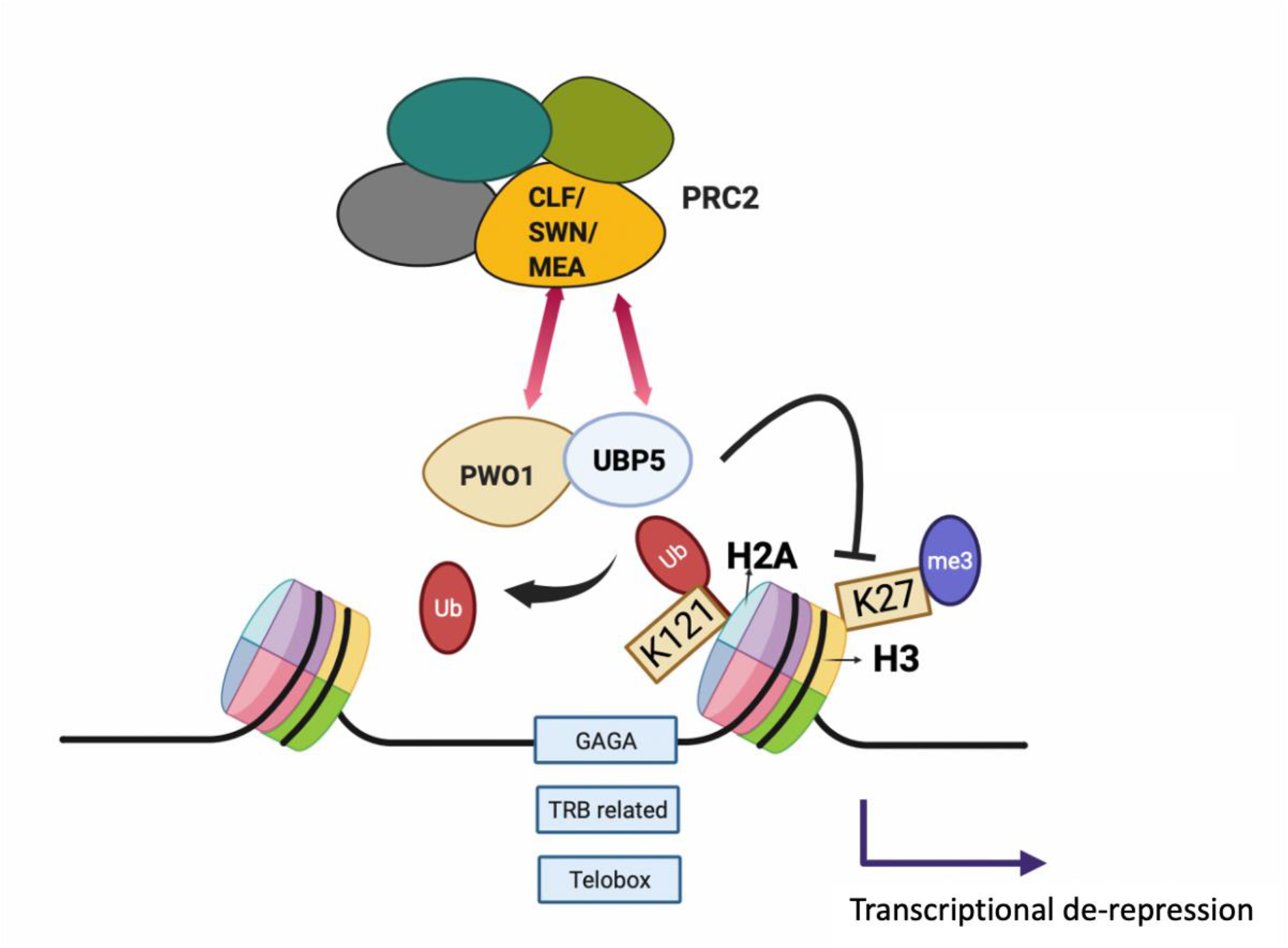
Working model for UBP5 function. UBP5 interacts with both PRC2 and PWO1 and its recruitment to chromatin associates with TRB- and PRC2-related *cis*-elements (light blue boxes). UBP5 acts as H2A deubiquitinase and prevents deposition of H3K27me3 leading to transcription de-repression. Figure is created using Biorender.

### UBP5 plays a role in transcriptional de-repression

Functional categorisation of UBP5 direct targets revealed that genes related to chromosome organisation, histone binding and chromatin binding were significantly over-represented (Supplementary Fig. 8A-C). In addition to UBP5 interaction with PWO1 and SWN chromatin factors, we identified its direct binding to several PRC2 subunit genes such as *CLF, EMF2, VERNALIZATION 2* (*VRN2*), *FERTILIZATION-INDEPENDENT ENDOSPERM* (*FIE*) and *MULTICOPY SUPPRESSOR OF IRA* 1 (*MSI1*) and PRC1 subunit encoding gene *B LYMPHOMA Mo-MLV INSERTION REGION ONE HOMOLOG* (*BMI1B*) (Supplementary list 6). H2Aub mark was also gained in these genes (Supplementary list 6), although for most of the genes we did not observe transcriptional changes in *ubp5*. On the other hand, GO analyses of UBP5 target genes that gained the H2Aub mark in *ubp5* revealed a significant over-representation of genes involved in response to DNA damage and repair (Supplementary Fig. 8D).

At the genome-wide level, UBP5 binding to chromatin typically occurs at the proximity of the transcription start site (TSS) and the start of the coding region (Supplementary Fig. 6A). Analyses of UBP5 binding peaks showed that majority of these sites correspond to protein coding genes, particularly exons and 5’UTRs that respectively correspond to ∼51% and ∼23% of the binding sites (Supplementary Fig. 9). Hence, we evaluated the impact of UBP5 in the transcriptional output of its target genes by integrating our ChIP-seq and RNA-seq data. We found a clear link between UBP5 gene binding and repression since 43% (207/478) of the genes downregulated in *ubp5* correspond to UBP5 targets gaining H2Aub in *ubp5*, whereas UBP5 is almost never found associated to upregulated genes (4/345 genes) (Fig. 4A-B). More generally, *ubp5* associated defects in transcription and H2Aub levels globally correlate (Fig. 4C-D), suggesting a role of UBP5 in relieving H2Aub-mediated repression, thereby promoting gene expression. Therefore, UBP5 seems to be predominantly involved in H2Aub erasure, which, at least for a set of its targets genes, results in transcriptional de-repression.

### UBP5-mediated H2A deubiquitination prevents deposition of H3K27me3

To explore whether UBP5 is targeted to chromatin in a sequence-specific manner, we analysed sequence motifs at UBP5 binding sites using MEME-ChIP ^28^ and identified a significant over-representation of GAGA, Telobox and Telobox-related motifs (Fig. 5A). Notably, GAGA elements recognised by transcription activators/repressors and Telobox motifs typically recognised by TRBs, are involved in recruiting PRC2 and TRBs together with PWOs form part of the PEAT complex ^17, 29, 30^. These results thus suggest the existence of sequence-specific mechanisms commonly recruiting UBP5, PWO proteins and PRC activity.

Therefore, to further unravel the relationship between UBP5 function and PRC2 activity, we analysed H3K27me3 bulk level by WB analysis and identified a 70% increase in its abundance in *ubp5* (Fig. 5B). We conducted ChIP-seq to further determine the genome-wide effects of UBP5 on H3K27me3. Our data showed a high overlap of H3K27me3 marked genes in Col-0 seedlings with previously published data (^31^; Supplementary Fig. 10A; Supplementary list 3). In addition, our genome-wide data showed that, at UBP5 target genes, H3K27me3 level was higher on average in *ubp5* (Fig. 5C). Notably, high H3K27me3 level was particularly pronounced at gene domains corresponding to UBP5 binding sites (Fig. 5C-D). Differential analysis of H3K27me3 marks revealed 2,587 H3K27me3 hyper-marked and 2,363 H3K27me3 depleted genes in *ubp5* (Supplementary list 7). Further analyses of ChIP-seq data based on differential analysis showed that in 602 genes the following conditions concurred: i) H3K27me3 and ii) H2Aub gained in *upb5*, and iii) directly bound by UBP5 (Fig. 5E and 5F; Supplementary Fig. 10B), indicating that UBP5 not only erases H2Aub but also affects H3K27me3 at multiple sites. In addition, our data revealed that only 3% of H3K27me3 depleted genes were UBP5 targets (Supplementary Fig. 10C), suggesting that UBP5 may not play a direct role in H3K27me3 maintenance at these genes and therefore these changes might likely result from indirect effects in the regulation of H3K27me3 writers’ or erasers’ activity. In agreement with a repressive role of H3K27me3 marking, average H3K27me3 levels in the gene body of *ubp5* downregulated genes was significantly higher than Col-0 levels, and there were no significant changes in the upregulated genes (Fig. 5G) and, similarly, we found a correlation between H3K27me3 and transcript levels in *ubp5* (Fig. 5H). Hence, UBP5 may de-repress such genes by preventing H3K27me3 enrichment.

To understand how both H2Aub and H3K27me3 dynamics affect the transcriptional levels of genes we focussed on the set of genes which gained H2Aub in *ubp5*. In this set of genes, we analysed the transcriptional levels of H3K27me3/H2Aub marked genes in both Col-0 and *ubp5* and found that in both background, genes that are exclusively marked by H2Aub are more highly expressed than genes with the two marks or only H3K27me3, as previously shown (Zhou *et al*., 2017). On the other hand, while in Col-0 plants there is a significant difference in transcriptional levels of H2Aub/H3K27me3 versus H3K27me3 marked genes, this difference is lost in *ubp5* with both categories showing similar repressive levels (Supplementary Fig. 10D). Hence, UBP5 may contribute to pose H2Aub/H3K27me3 marked genes in a more responsive chromatin structure. Overall, we thus conclude that in the subset of 602 genes, UBP5-mediated H2Aub deubiquitination prevents the deposition of H3K27me3 mark leading to a de-repressed chromatin environment (Fig. 6).

## Discussion

PRC2 interactors play a key role in regulating its molecular activities and recruitment to chromatin ^7^. For instance, we previously showed that PWO1 may mediate in providing PRC2 with the right chromatin environment to methylate H3 ^14^. In addition, PWO1 was proposed to form part of the PEAT complex mediating silencing ^15^. Therefore, unravelling the protein interactors associated with epigenetic pathways can provide important clues to understand their possible crosstalk and activities. Here, we have demonstrated that UBP5 is a novel interactor of PWO1 and PRC2. UBP5 was also identified co-immunoprecipitating with all main components of PEAT ^15^. Most deubiquitinases may require to be in multi-subunit complexes to be enzymatically active, as it has been shown for H2A deubiquitinases Myb-like SWIRM (2A-DUB) and USP22 in human cells ^32, 33, 34^ or H2B deubiquitinase UBP22/USP22 in plants and other eukaryotes ^35^. On the other hand, UBP5 binding sites are enriched in *Telobox* and other telomeric related motifs that have been previously involved in PRC2 recruitment by TRBs at genes ^17^ and telomeric regions ^36^. TRBs are also one of the components of the PEAT complex ^15^. Therefore, a plausible hypothesis is that sequence-specific UBP5 chromatin association is, at least in part, driven by its interaction with PWO1 in the frame of the PEAT complex. Future analyses to identify UBP5 protein network will also help to confirm whether UBP5 associates to TRBs and/or other PEAT subunits. Furthermore, whether UBP5 forms a stable complex or a more dynamic protein network with its interactors and whether its activities depend on, or are independent of, these interactions will be important questions to address.

UBP5 belongs to the UBP family, which is part of the conserved DUB superfamily. Several DUBs are involved in the regulation of chromatin and some of them especially in H2A deubiquitination ^11^. For instance, Drosophila protein Calypso as well as its corresponding ortholog in humans, the tumour suppressor BRCA-1-associated protein 1 (BAP1), form part of a PR-DUB complex able to remove the H2AK119ub1 mark. Intriguingly, PR-DUB has been described as a type of PRC despite its opposite activity to PRC1. Therefore, it seems that a dynamic ubiquitination/deubiquitination counterbalance is key for maintaining PRCs’ activities and proper H2A ubiquitination levels over the genome ^37, 38, 39^. Phylogenetic analyses confirmed that there are three proteases in Arabidopsis, UCH1-3, that belongs to the same family as Calypso/BAP1; however, it is unknown if any of them have conserved a similar function in plants ^11^. Indeed, UCH1-3 have recently been related with the control of the circadian clock oscillation under high temperatures ^40^ and previously with the response to auxins during development ^41^, but no data link these proteins to chromatin regulation so far. The only proteins that have been related to H2A deubiquitination in Arabidopsis are the closely related UBP12 and 13 proteins, which were identified interacting with LHP1 ^26^, a protein that may act as an accessory protein in both PRC2 and PRC1 ^7^. UBP12 was shown to be involved in the repression of a subset of PRC2 targets mediating H3K27me3 deposition and to be actively involved in H2A deubiquitination ^26^. UBP12/13-mediated H2Aub removal prevents loss of H3K27me3 and therefore these proteins may be involved in stable PRC2-mediated repression ^13^. In contrast, our data indicate a role of UBP5 in preventing H3K27me3 gain at specific loci (Fig. 6). Moreover, the genes that are regulated by UBP12/13 (i.e. H2Aub gained genes in *ubp12/13*) and UBP5 direct targets show little overlap (Supplementary Fig. 11A), suggesting that they act through independent mechanisms or at different genome domains. However, as UBP12/13 direct target genes have not been described so far, this conclusion needs to be cautiously considered as indirect results in *ubp12/13* epigenomic data cannot be discarded ^13^.

UBP12/13 are the closest Arabidopsis orthologs to UBIQUITIN SPECIFIC PROTEASE 7 (USP7) in animals ^11^. In Drosophila, USP7 has been involved in the regulation of PcG targets and in gene silencing through heterochromatin formation, which seems to play a key role in genome stability ^42^. In addition, studies using cancer cell lines demonstrated that USP7 directly interacts and stabilises EZH2, the HMT of PRC2 ^43^, and PRC1.1, one of the human PRC1 complexes ^44^, indicating another scenario for the activities of the USP7 like proteins ^45^. On the other hand, UBP5 closest human orthologs are USP4, USP11 and USP15 ^11^. Among them, *USP11* has been described as an oncogene that regulates cell cycle and cancer progression through DNA repair. USP11 acts in both H2AK119 and H2BK120 deubiquitination as part of the nucleosome remodelling and deacetylase (NuRD) complex and specifically deubiquitinates γH2AX, which is key in homologous recombination ^46^. Our ChIP-seq profiling in seedlings identified that UBP5 is required for H2Aub deubiquitination at a majority of PRC1-regulated Arabidopsis genes, and, considering *ubp5* phenotypes, UBP5 may have additional effects on H2Aub epigenome at other developmental stages. H2Aub ChIP-seq profile also points to a dual role of UBP5 deubiquitination activity. In ∼40% of genes showing a H2Aub gain in *ubp5*, UBP5 acts to maintain a certain level of H2Aub in the plant; while, in ∼60% of this set of genes, UBP5 fully erases this histone mark. Overall, these results indicate that UBP5 acts in *cis* to maintain the right H2Aub level at target genes with two possible scenarios for each locus: this modification is either 1) erased by UBP5 in most cells and therefore not detected in Col-0 plants but only in *ubp5* (i.e. *de novo* marked genes) or 2) stably present in Col-0 seedlings but removed by UBP5 only in certain genome copies or in certain cells (i.e. H2Aub hyper-marked genes). Further studies will be required to fully understand how UBP5 discerns between these different scenarios.

Therefore, our results point to a conservation between Arabidopsis UBP5 and human USP11 activities as H2A deubiquitinases. Whether UBP5 may have additional roles in DNA repair as USP11 will require further investigation, but the fact that many H2Aub-enriched UBP5 target genes are related with DNA damage and binding supports this possibility. As our H2Aub ChIP-seq data was obtained for the bulk of this histone modification, we cannot rule out that these epigenomic data in fact reflects the ubiquitination status of specific H2A variants. Thus, it will be very interesting to test if UBP5 differentially affects the post-translational modifications of H2A variants, such it has been shown for H2AX deubiquitination by USP11 ^46^. Another exciting possibility to explore will be the deubiquitination of the H2A.Z histone variant, which ubiquitination is mediated by PRC1 to induce PRC2-independent transcriptional repression ^47^. The possibility that UBP5 mediates H2A.Z deubiquitination is supported by the remarkable overlap between H2A.Z marked genes and UBP5 direct targets that gained H2Aub in *ubp5* (Supplementary Fig. 11B), opening future venues to further understanding UBP5 activities.

Mirroring the meta-gene pattern of H2Aub in Arabidopsis (^31^; Fig. 3D), UBP5 predominantly binds to chromatin in the vicinity of TSSs and at the start of protein coding regions. Furthermore, our transcriptional analyses show that UBP5 target genes tend to be downregulated in the *ubp5* mutant. These results point to UBP5 acting as a transcriptional activator, as shown for H2A deubiquitination in animals ^48^. As UBP5 acts in histone deubiquitination, we favour the possibility of its active role in promoting transcriptional de-repression through the erasure of H2Aub as it has been proposed for other erasers (e.g. histone demethylases ^49^). However, gain of H2Aub in *ubp5* is not always synonymous of changes in transcription in a comparable way as accessible chromatin is not always leading to activation ^50^.

Our expression analyses in *ubp5* also indicate that stress responsive genes are among the most affected. Notably, it has been proposed that H2Aub is involved in creating a repressive but reactive chromatin environment ^50^ and, thus, UBP5 may be a key factor in positively regulating the chromatin of genes that need to respond to specific environmental signals. Indeed, the combined analyses of the transcriptomic and epigenomic data in WT versus *ubp5* showed that, while having only H3K27me3 is more repressive than being marked by H2Aub and H3K27me3, both in previous ^31^ and in our data, this difference is lost in *ubp5*. This may suggest that UBP5 is essential to keep H2Aub under a certain threshold that helps H2Aub/H3K27me3 marked genes to be more reactive. On the other hand, PWO1 was proposed to mediate PRC2-related repression of stress responsive genes ^18, 51^. A possible scenario is that UBP5-PWO1 antagonistic activities, respectively as activator and repressor, create a bistable and more responsive chromatin.

In line with the UBP5-PRC2 protein interaction identified here, UBP5 influences H3K27me3 levels at a majority of H3K27me3-marked genes (4,950 out of 7,600 genes), ∼20% of them corresponding to direct target sites at the seedling stage (1,013 genes). For these genes, deposition of H2Aub plausibly precedes H3K27 trimethylation on the same nucleosome, as suggested for several PRC1/PRC2 target genes ^52^, and hence UBP5-mediated deubiquitination will prevent H3K27me3 deposition by PRC2 (Fig. 6), probably making chromatin more accessible in these loci. Our proposed functional model also fits well with evolutionary results linking the deposition of H3K27me3 to the ubiquitination of H2A in *Marchantia polymorpha* ^53^. Despite all our results leading to an UBP5-PRC2 interaction, we should not forget that many UBP5 target genes that are enriched in H2Aub do not gain H3K27me3, indicating that UBP5 plays PRC2-independent functions. This opens further fascinating questions about UBP5 alternative activities in controlling chromatin accessibility that we look forward to answering in future studies.

## Materials and Methods

### Plant Materials and Cultivation conditions

All *Arabidopsis thaliana* (Arabidopsis) lines used in this study were in the Columbia-0 (Col-0) ecotype background. For the generation of *ubp5* CRISPR-Cas9 mutant, double guide system of Cas9-directed mutagenesis was performed as described by ^54^ to delete a fragment size of 3,361 bp from *UBP5* gDNA sequence (Supplementary Fig. 2A). sgRNAs were designed using CRISPR-P tool ^55^. The P3-Cas9-mCherry vector for generating the *ubp5* line was kindly provided by Charles Spillane’s lab ^54^. Deletion of the genomic fragment from *UBP5* was confirmed using Sanger sequencing (LGC genomics, Germany). Transgenic plants were developed by *Agrobacterium*-mediated gene transformation with floral dip method ^56^. For genotyping, DNA extraction was done based on ^57^. Oligonucleotide primers used for CRISPR-Cas9 mutagenesis and genotyping are indicated in Supplementary Table 1. For the *UBP5pro::UBP5-GFP;ubp5* line, a 1,708-kb-upstream fragment and gene-body regions of *UBP5* without stop codon were amplified from genomic DNA of Col-0 with GW-compatible primers (Supplementary Table 1). *gUBP5* was fused with a C-terminal GFP sequence in the (pGKGWG) vector ^58^.

Sterilised seeds were sown on Murashige & Skoog medium (MS Base) supplemented with 1% Sucrose, 0.1% MES, 0.8% agar with pH adjusted to 5.6, stratified at 4 °C for three days and placed to Percival tissue culture cabinet under a 16:8 h light: dark (21°C/18°C) regime until they were transferred to soil. Arabidopsis plants were grown on pots containing compost, vermiculite and perlite (5:1:1 proportion) with the same photoperiod under fluorescent lamps at 200 μmol m^−2^ s^−1^. For hypocotyl and root length measurements, Col-0 and *ubp5* seeds were sown on MS medium, and the plates were placed vertically in the growth chamber in LD conditions. Photographs were taken at the end of 10 days, hypocotyl and root length were measured using the Fiji image processing software.

### Yeast two hybrid assay

For yeast two hybrid assays, untransformed *Saccharomyces cerevisiae* AH109 cultures were grown at 28 °C, on solid or liquid Yeast Peptone Dextrose (YPD) media supplemented with adenine (80 mg/L). The *S. cerevisiae* AH109 competent cells were obtained as previously described ^59^. For Yeast two hybrid (Y2H) experiments, yeast were co-transformed using a heat shock method at 42°C for 30 min ^60^. For plating, 3 μl of culture were plated at the same concentration on drop-out media (minimal medium) in the absence of leucine and tryptophan (SD-L-W) or more restrictive media without histidine (SD-L-W-H) in serial dilutions. Yeast growth was analysed after 3 to 4 days growing at 28°C. Both bait and prey empty vectors were used as negative controls.

### Co-immunoprecipitation assay

Modified versions of pMDC7 carrying the GFP or mCherry tags ^61^ were used to insert the coding sequence of *UBP5* and *SWNΔSET* via Gateway cloning (Invitrogen). Vectors were transformed in *Agrobacterium tumefaciens* (Agrobacterium) GV3101 pMP90. For transient expression assays, the abaxial sides of leaves of 4/5-week-old *Nicotiana benthamiana* plants were infiltrated with transformed Agrobacterium cell culture suspension in log phase growth. Expression was induced by spraying 20 μM β-estradiol in 0.1% Tween onto infiltrated leaves 48 to 72 h after Agrobacterium infiltration. Fluorescence was monitored in leaf epidermis cells after a short induction period (4–6 h when fluorescence was visible) using an Olympus BX51 epifluorescence microscope. After 6 h from the second induction of β-estradiol, the samples were frozen in liquid N_2_. The samples were ground in a liquid N_2_ pre-cooled mortar followed by 20 min at 4°C in a shaker in 10 ml of protein extraction buffer (10% glycerol, 150 mM NaCl, 2.5 mM EDTA, 20 mM Tris-HCl pH 8, 1% Triton and Complete® EDTA-free protease inhibitor cocktail (1 tablet/50 ml; Roche)). After resuspension, samples were filtered through two Miracloth (Calbiochem®) layers and centrifuge at 4°C 15 min 4,000 rpm. After centrifugation, the supernatants were transferred to a new 15 mL tube, and the extracts were taken, mixed with 3X Laemmli buffer (0.3 M Tris-HCl (pH 6.8); 10 % (w/v) SDS; 30 % (v/v) glycerol; 0.6 M DTT; 0.01% (w/v) bromophenol blue) and heated at 95°C for 5 min. Co-IPs were carried out by incubating the samples with 30 μL of protein A agarose bead slurry for 4h at 4°C in a rotating wheel and with anti-mCherry (Takara 632496) of 1:1000 dilution. After 4 h incubation, a centrifugation at 4°C at 500 g for 2 min was carried out to precipitate the beads. The beads were washed 3 times with protein extraction buffer, resuspended in 3× Laemmli buffer and denatured at 95°C for 10 min. Proteins were loaded in 10% SDS-PAGE gels and transferred to a PVDF membrane. Membranes were developed with anti-GFP (Roche 11814460001).

### Subnuclear Localisation and FRET assay

For subnuclear localization in *N. benthamiana*, estradiol-inducible pMDC7-derivatives plasmid vectors containing our coding sequences were transformed into Agrobacterium (GV3101 PMP90 strain with p19 silencing suppressor plasmid). FRET assay was performed as described in ^18^. Images were captured by confocal microscopy on a LSM780 (Zeiss) or SP8 (Leica).

### Histone extraction and Western Blot

Nuclei were extracted from 1.5 g of 12 days after germination (DAG) seedlings using the nuclei extraction buffer (0.4 M Sucrose, 10mM Tris-HCl pH 8.0, 5mM β-Mercaptoethanol, 10mM MgCl_2_, 0.1mM PMSF). Extracted nuclei were treated overnight with 0.4 N H_2_SO_4_ to obtain a histone-enriched extract. The extracted proteins were precipitated with 33% trichloroacetic acid and then washed 3 times with acetone, air-dried, and re-suspended in 100 μL 3X Laemmli buffer. The samples were boiled for 10 min, separated on 15% sodium dodecyl sulfate-polyacrylamide electrophoresis gels and transferred to a polyvinylidene difluoride membrane (Immobilon-P Transfer membrane, Millipore) by wet blotting in transfer buffer (25 mM Tris– HCl, 192 mM glycine, and 10% methanol). Primary and secondary antibodies used were anti-H2Aub antibody (Cell Signalling Technology D27C4), anti-H2A antibody (Active Motif 91325), anti-H3K27me3 antibody (Millipore 07-449), anti-H3 (Abcam ab1791), anti-mouse IgG (H+L) HRP conjugated (Chemicon International AP308P) and Anti-Rabbit IgG (whole molecule)–Peroxidase (Sigma Aldrich A9169). Chemiluminescence detection was done with SuperSignal West Pico or Femto (Thermo Fischer Scientific) following the manufacturer’s instructions.

### ChIP-qPCR, ChIP-seq and Data analyses

Chromatin immunoprecipitations (ChIP) were carried out using 12-DAG seedlings as described previously ^25^. Chromatin was extracted from formaldehyde fixed tissue and fragmented using a Bioruptor® Pico (Diagenode) in fragments of 200–500 bp. Antibodies used for ChIP-qPCR in this study were H3K27me3 (Millipore 07-449) and H2Aub (Cell Signalling Technology D27C4). 30 μl/sample of Protein A Dynabeads (10002D) were used for preclearing before IP. The IP was performed with 60 μl/sample of Protein A Dynabeads and 5 μl of antibodies in the ChIP dilution buffer at 4°C overnight. Following IP, chromatin was washed with four different wash buffers-Low Salt, High salt, LiCl and TE wash buffer sequentially. Then, the chromatin was eluted and crosslinking was reversed overnight at 65°C. After IP, DNA was eluted and purified using ultrapure phenol:chloroform:isoamyl alcohol (25:24:1) pH 8.05 followed by ethanol precipitation. Input DNA was diluted to 1:10, and 1 μl of IP DNA was used for quantitative PCR (qPCR). ChIP-qPCRs were carried out in a CFX96TM Real-Time PCR Detection System (Bio Rad) using TakyonTM No Rox SYBR MasterMix dTTP Blue (Eurogentec). Oligonucleotide primers used for ChIP-qPCR are listed in Supplementary Table 1.

For ChIP-seq experiments, chromatin extraction and immunoprecipitation of histones were done as previously described ^25^ in three biological replicates for H2Aub and two biological replicates for H3K27me3 at 12-DAG old Col-0 and *ubp5* seedlings grown under LD conditions. Two IPs were carried out for each biological replicate using 100 μg of chromatin, quantified using Pierce BiCinchoninic Acid (BCA) assay kit (Thermo Fisher Scientific). After IP, DNA was eluted and purified. Library preparation and paired end sequencing was performed using DNA Nanoballs (DNB™) sequencing technology from BGI (Sequencing method: DNBSEQ-G400_PE100). Reads were mapped using STAR v2.7.8a ^62^ onto TAIR10 Arabidopsis with parameters align intron max as 1 and align ends type as EndToEnd. The organelle genomes were excluded from the mapped reads. Duplicated reads were removed using Picard tool MarkDuplicates option. Only uniquely mapped reads were retained for further analysis. Marked peaks for each IP were obtained using MACS3 ^63^ with parameters broad peak and q value cut off as 0.05. Browser tracks were obtained using the bamCoverage function by scaling with the parameter --normalizeUsing RPGC. Tracks were visualised using IGV v2.12.3 ^64^. Bedtools Utility Intersect ^65^ was used to intersect the MACS3 peaks obtained from the biological replicates. The resulting peaks from the biological replicates were merged and annotated with TAIR10 gene coordinates. To determine gain or depletion of H2Aub or H3K27me3 marks, the number of reads mapping into the peak coordinates was calculated using Bedtools Utility Multicov and the peaks from all samples were grouped by gene-ID to obtain unique peak coordinates per marked gene using Bedtools Utility Groupby v2.26.0 ^65^. Differential enrichment of respective marks between samples were done using DESeq2 analysis ^66^. The comparison between biological replicates of H2Aub and H3K27me3 are shown in Supplementary Fig. 12 and 13.

### UBP5-GFP ChIP-seq and data analyses

UBP5-GFP ChIP was performed with *UBP5pro::UBP5-GFP;ubp5* line using a double crosslinking protocol as described ^67^. Two biological replicates with 2 g each from 12-DAG seedlings were ground in liquid N_2_ to fine powder and resuspended in nuclei isolation buffer (60 mM HEPES pH 8.0, 1 M Sucrose, 5 mM KCl, 5 mM MgCl_2_, 5 mM EDTA, 0.6% Triton X-100, 0.4 mM PMSF, pepstatin and complete protease inhibitors (Roche). Then, the samples were cross-linked with 25 mM ethylene glycol bis succinimidyl succinate (EGS) by rotating for 20 min and with 1% formaldehyde by rotating for 10 min. The crosslinking of samples was stopped by 2M glycine for 10 min at room temperature. The chromatin was isolated and sheared into 200–500 bp fragments by sonication. For IP, the sonicated chromatin was incubated with 20 μl of anti-GFP antibody (Thermo Fisher #A11122) overnight at 4°C while gentle rotating. Followed by IP, eluted and purified DNA of two independent biological replicates along with input control without antibody was used for library preparation and paired end sequencing was performed using DNB™ sequencing technology from BGI.

For UBP5-GFP ChIP-seq data analysis, Raw data with adapter sequences or low-quality sequences was filtered using SOAPnuke software (BGI). The reads were mapped to the Arabidopsis genome (TAIR10) using Bowtie2 2.4.5 ^68^ with default parameters. Only uniquely mapped reads were retained for further analysis. Peaks were called using MACS3 ^63^. The peaks were converted to bigwig files using deepTools ^69^. bamCoverage was done using RPGC normalisation. The intersections of common peaks between two biological replicates with FDR < 0.01 was obtained using Bedtools Utility Intersect v2.30.0 ^65^. The oligonucleotide primers used to confirm few UBP5-target genes using ChIP-qPCR are listed in the Supplementary Table 1. Comparison between ChIP-seq replicates were shown in Supplementary Fig. 13

For DNA motifs analyses, we considered −500 bp to +250 bp from TSS for the UBP5 target genes using ‘getfasta’ function. We searched for enriched DNA motifs using the fasta file as a input for MEME-ChIP ^28^ with discriminative mode using the negative control sequences wherein UBP5 targeting regions were removed.

### RNA isolation, quantitative RT PCR

Total RNA was isolated from 12-DAG seedlings (Col and *ubp5*) using E.Z.N.A. Plant RNA Kit (OMEGA biotek) following manufacturer instructions. The RNA concentration was determined using the Nanophotometer (IMPLEN). RNA was examined by electrophoresis on a 1.2% agarose gel. For cDNA synthesis, RNA samples were subjected to DNAse treatment and cDNA synthesis was performed using (Thermo Scientific). Quantitative real time PCR (qRT-PCR) was performed in a CFX96™ Real-Time PCR Detection System (Bio Rad) using TakyonTM No Rox SYBR MasterMix dTTP Blue (Eurogentec). Expression levels were normalised to the reference genes *At5G25760* and *At4G34270* ^70^. Relative enrichment was calculated using the 2^-ΔΔCT^ method ^71^

### RNA-seq library preparation, sequencing and bioinformatics

For RNA-seq, RNA was extracted from 12-DAG seedlings with four biological replicates for each background (Col-0 and *ubp5*). Library preparation and RNA-seq was performed according to the protocol described recently ^72^. 500 ng DNase-treated RNA was used for reverse transcription with 50 mM different barcoded oligo(dT) primers and SuperScript III. Each reaction was pooled, pools were Ampure purified (1.5x beads to sample volumes) and then eluted. Second-strand synthesis was carried out using nick translation protocol (Krzyszton et al. 2022). Tagmentation reaction ^73^ was performed out using recovered dsDNA sample incubated with homemade Tn5 enzyme in a freshly prepared 2x buffer (20 mM Tris-HCl pH 7.5, 20 mM MgCl2, 50% DMF). Illumina indexing PCR was performed using the tagmented DNA. Libraries were sequenced on Illumina NextSeq 500 system using the paired-end mode to obtain 21 nt R1 (contain barcode and Unique Molecular Identifier (UMI)) and 55 nt R2 (contain mRNA sequences).

After quality control using fastqc, reads R1 and R2 were processed separately. In our oligo(dT) primers two parts of UMI are split by barcode sequence, therefore we transformed read R1 fastq file using awk command. Read R2 was trimmed to remove potential contamination with poly(A) tail using BRBseqTools v 1.6 Trim ^74^. Reads were mapped using STAR v 2.7.8a ^62^ to TAIR 10 genome with Araport11 genome annotation. Finally, the count matrix for each library and each gene was obtained using BRBseqTools (v 1.6) CreateDGEMatrix ^74^ with parameters *-p UB -UMI 14 -s yes*, using Araport11 genome annotation and a list of barcodes. The differential gene expression analysis was done using the DESeq2 ^75^. Further, the genes were filtered based on log2 fold-change of ±1 and an adjusted p-value of less than 0.05 and categorised as upregulated, downregulated, and unaltered genes. GO enrichment analysis was performed in different gene set using ShinyGO tool ^76^.

## Authorship contributions

JG and SF conceptualised the experiment approach and designed the methodology; JG, EM, LW and JL performed the experiments; JG, MG, AF, MK, FB and CB performed the genomic data curation, analysis, and visualisation; JG and SF wrote the original manuscript; JG, FB, CB, SS, DS, and SF contributed to the interpretation of results; all the authors contributed to manuscript revision and approved the final manuscript.

## Acknowledgements

SF acknowledges support from the College of Science and Engineering (University of Galway). SF is grateful to Jennifer Siobal and Ronan Halton for technical support.

