## Supplementary Figures for "The UBP5 histone H2A deubiquitinase counteracts PRC2-mediated repression to regulate Arabidopsis development and stress responses"

**James Godwin et al**

### Supplementary Information

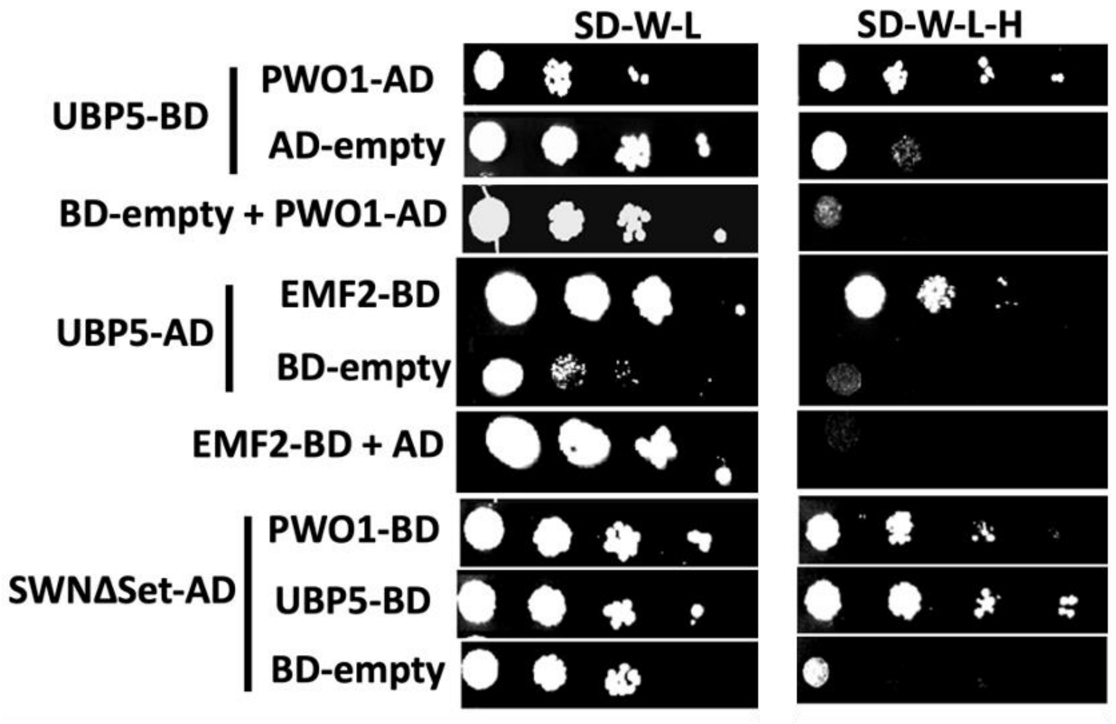


**Supplementary Figure 1. UBP5 interacts with other PRC2 protein.** Y2H analyses showing the interaction of UBP5 with EMF2 (PRC2 component). PWO1-SWNΔSet and PWO1-UBP5 interactions are shown as positive controls. Yeast cells containing the different construct combinations on selective medium for plasmids (-LW; -leucine, tryptophan) or for reporter gene activation (-LWAH; -leucine, tryptophan, adenine, histidine). Serial solutions were used. BD, GAL4-DNA binding fusion; AD, GAL4-DNA activation domain fusion. SWNΔSET, SWN construct lacking the SET domain.

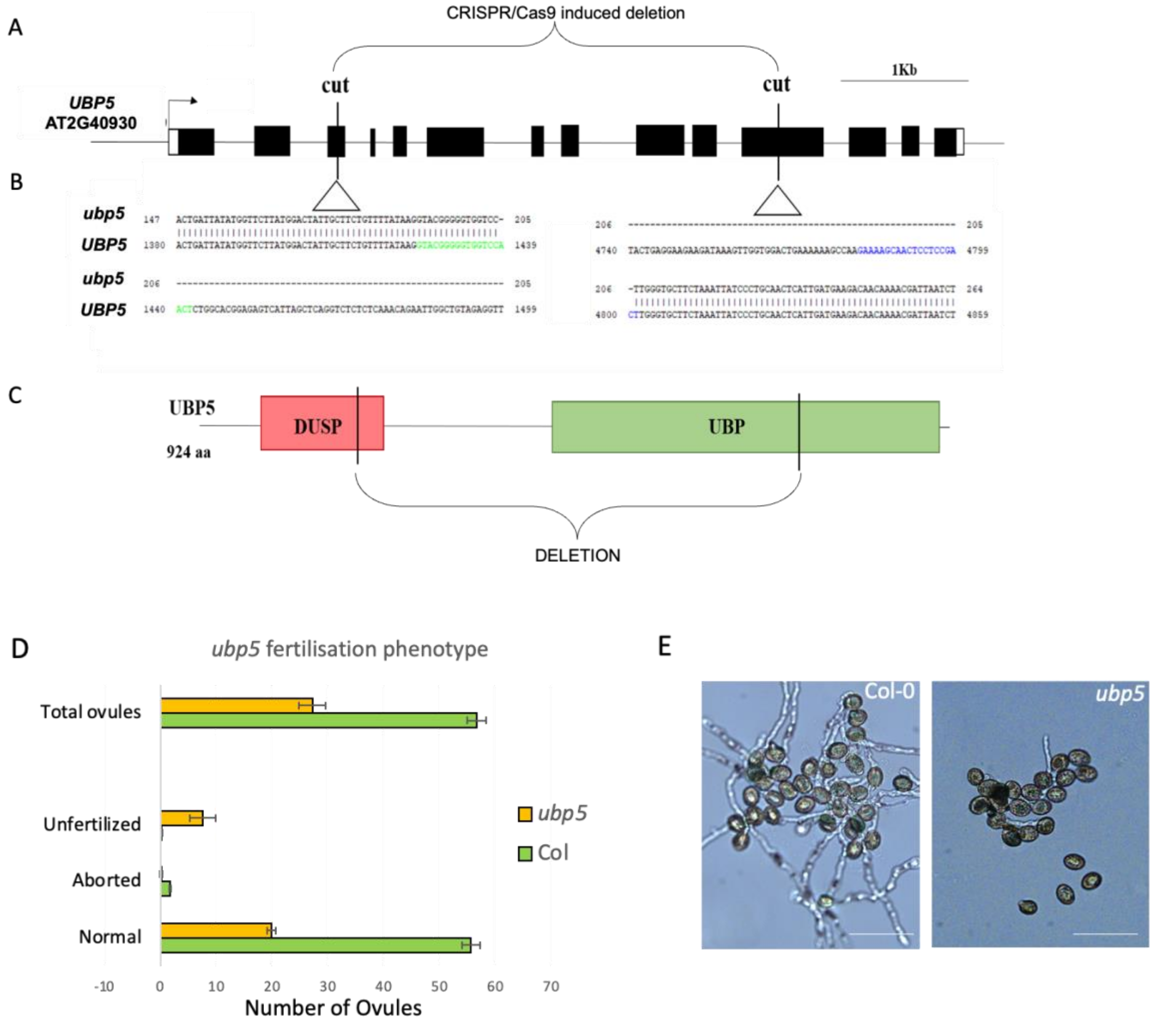


**Supplementary Figure 2. *ubp5* is a CRISPR/Cas9 induced deletion mutant with fertilisation defects.** A, Schematic representation of the *UBP5* (At2G40930) gene indicating the genomic region that was deleted by CRISPR/Cas9 in *ubp5* (black boxes, exons; white boxes, UTRs). B, Confirmation of the borders of the deletion in *ubp5* by sequencing. sgRNA regions marked respectively in green and blue. C, Schematic representation of the UBP5 protein indicating the deleted region in the protein encoded by the *ubp5* allele. D, Fertilisation analyses of *ubp5*. Fertilised versus normal ovules, aborted and unfertilised ovules in siliques were counted under the microscope (n = 15 siliques were used for Col-0 and *ubp5)*. Error bars represent the standard deviation (SD). *E, In vitro* pollen germination assays showing *ubp5* pollen germination defects. Pollen collected from Col-0 and *ubp5* plants. Scale bar 100 µm.

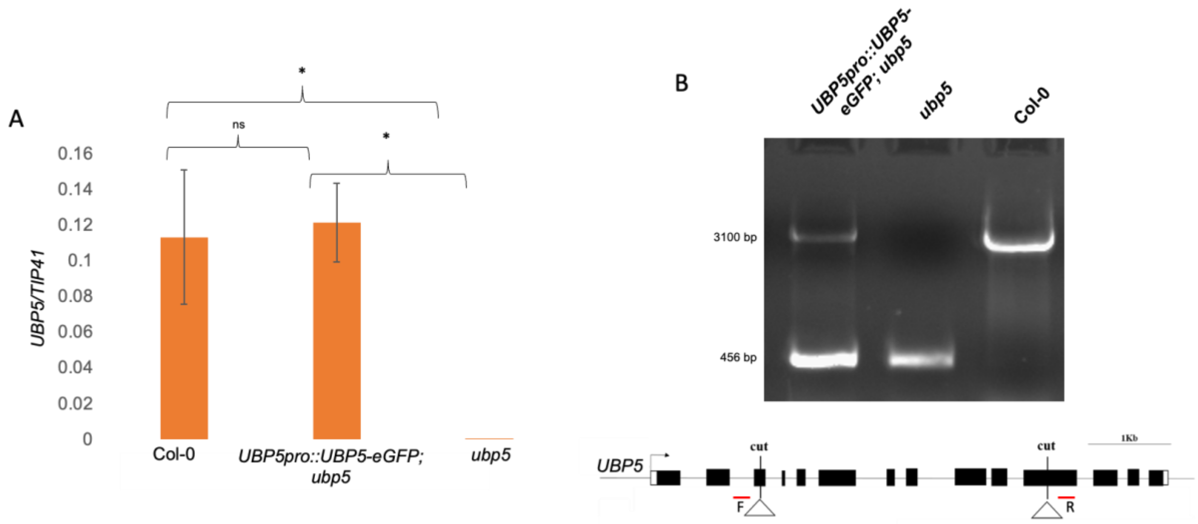


**Supplementary Figure 3. Complementation of the *ubp5* mutant by *UBP5pro::UBP5-eGFP*.** A, RT-qPCR analysis of *UBP5* expression in Col-0, *UBP5pro::UBP5-eGFP;ubp5* and *ubp5*. Error bars indicate SD of three biological replicates, grown and harvested independently. Significance differences indicated using Student's t-test represented by $p<0.05$ (*), ns represents non-significant. B, Genotyping results using forward (F) and reverse (R) primers showing that the complementing line *UBP5pro::UBP5-eGFP;ubp5* contains both wild type (upper band) and *ubp5* (lower band) alleles.

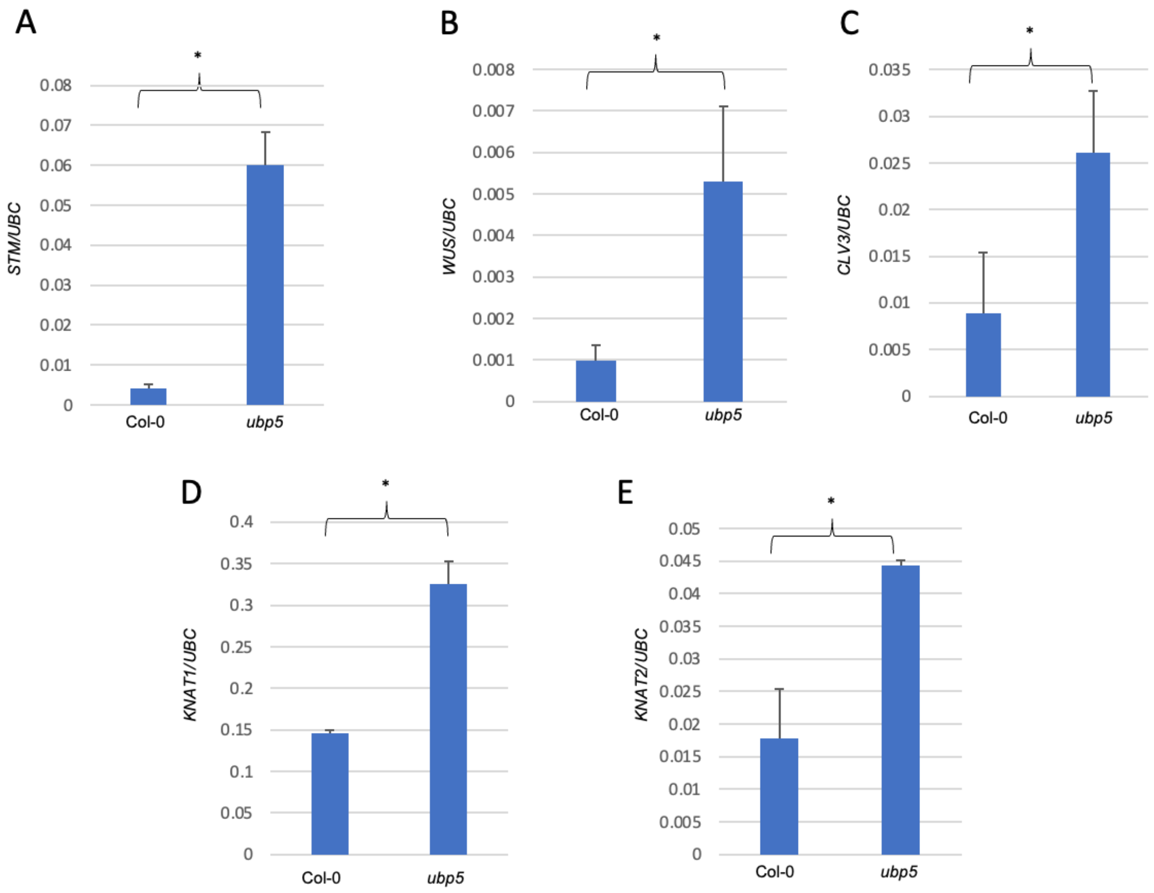


**Supplementary Figure 4. Expression analyses of PRC2 developmental target genes in *ubp5*.** A-E, RT-qPCR analyses showing that the expression of several shoot apical meristem (SAM) genes are affected in *ubp5:* A, *SHOOTMERISTEMLESS* (*STM*)*;* B, *WUSCHEL* (*WUS*)*;* C, *CLAVATA 3* (*CLV3*)*;* D, *KNOTTED-LIKE FROM ARABIDOPSIS THALIANA* (*KNAT1*)*;* E, *KNAT2*. Error bars indicate SD of three biological replicates, grown and harvested independently. Asterisks indicate significant differences calculated using Student's t-test (* p<0.05, ns – non-significant).

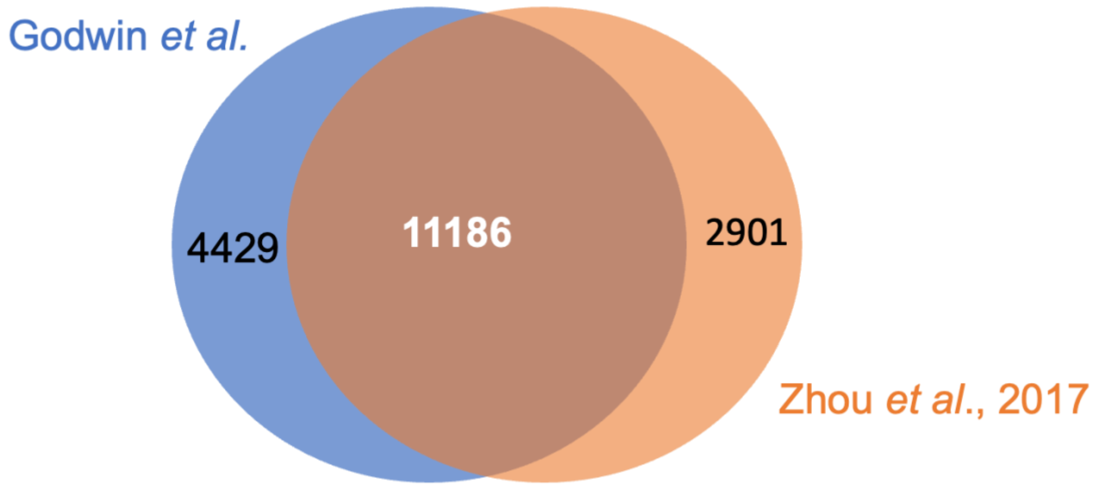


**Supplementary Figure 5. H2Aub marked genes between different datasets in Col-0 seedlings.** Venn diagram representing the high overlap of H2Aub marked genes in Col-0 between our dataset and Zhou et al., 2017 dataset.

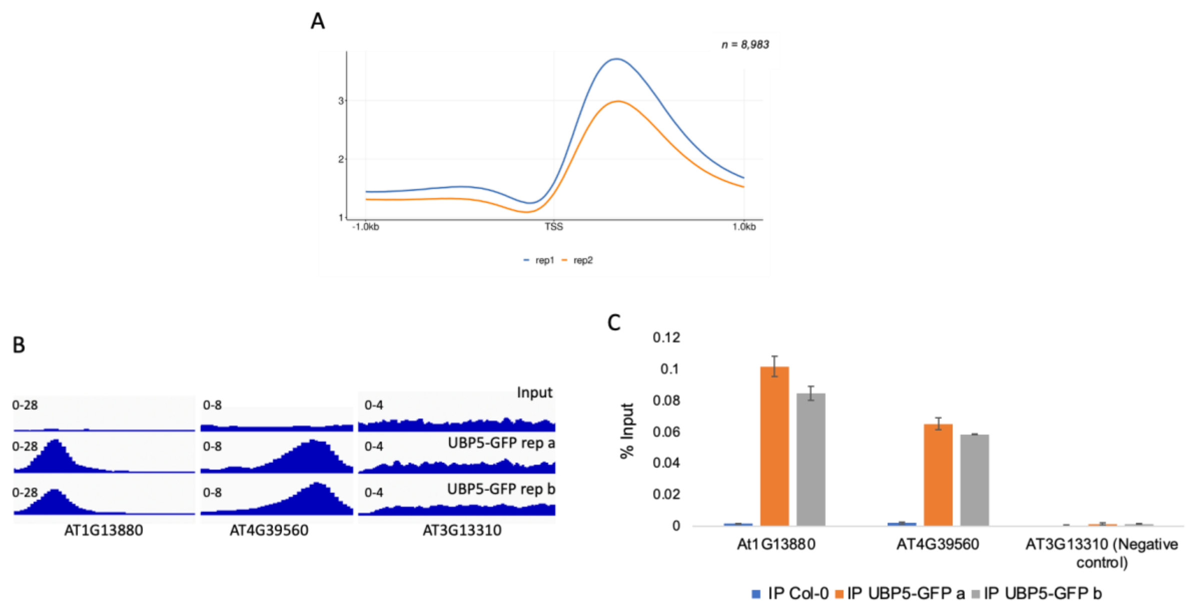


**Supplementary Figure 6. UBP5 enrichment at representative loci**. A, Metagene plot showing that individual biological replicates of UBP5 ChIP-seq. B, IGV browser view of UBP5 peaks from immunoprecipitated samples of *UBP5pro::UBP5-eGFP;ubp5* (UBP5-GFP) from two biological independent replicates a and b compared to input. C, ChIP-qPCR results for UBP5 target loci AT1G13880 and AT4G39560 in biological replicates a and b measured as % of input. AT3G13310 was selected as a negative control where no peak was observed in the IGV browser. Error bars indicate SD of two technical replicates.

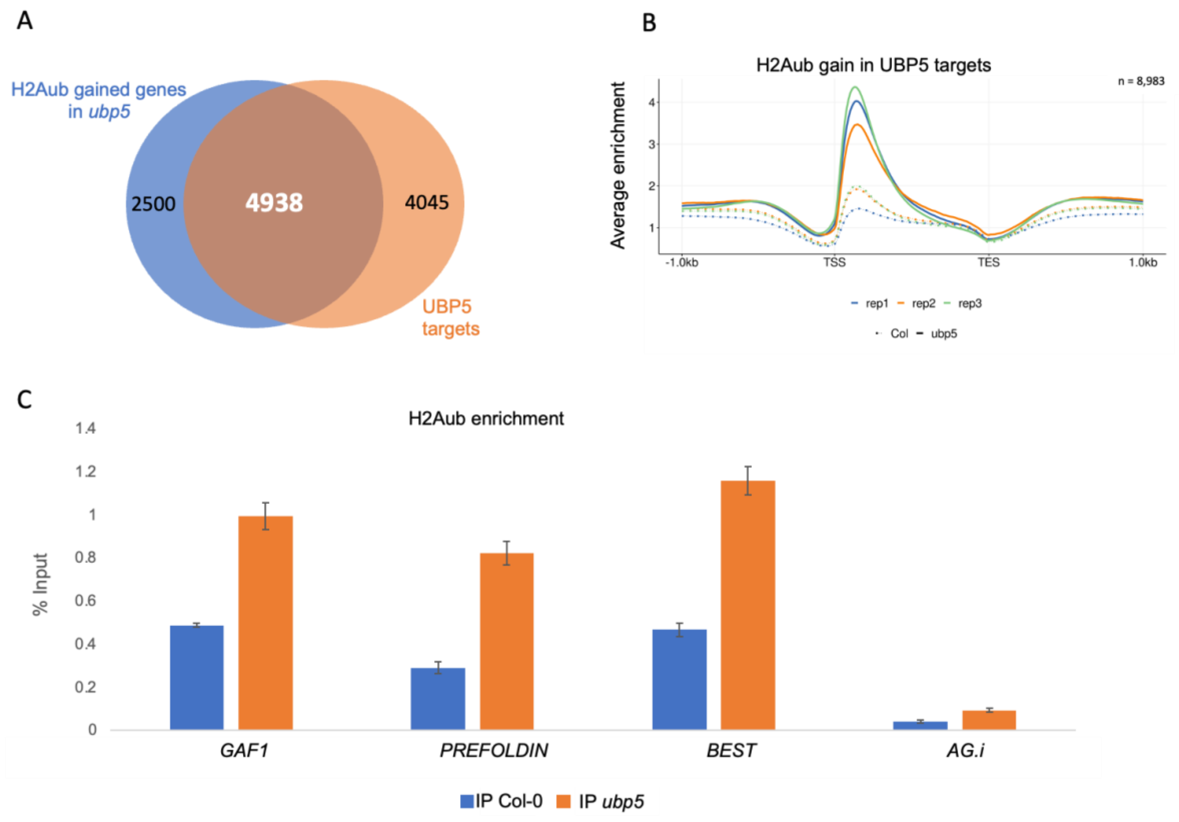


**Supplementary Figure 7. UBP5 binding overlaps with the gain on H2Aub in *ubp5*.** A, Venn diagram showing the overlap of genes between all H2Aub gained genes in *ubp5* based on DEseq2 analysis (FDR<0.05) and UBP5 direct target genes. B, Metagene plot of H2Aub distribution in the UBP5 targets for all three Col-0 and *ubp5* biological replicates. *C*, ChIP-qPCR validation of H2Aub hyper-marking in *ubp5*. Selected loci (*GAI ASSOCIATED FACTOR 1* (*GAF1* - AT5G59980), *PREFOLDIN* - AT1G03760 and *BESTROPHIN-LIKE PROTEIN* (*BEST* - AT3G61320)) were among the 207 UBP5 targets downregulated in *ubp5*. An intergenic region (*AG*.i) which is not H2Aub enriched was selected as negative control. ChIP-qPCR results shown as % of input DNA. Error bars indicate SD of three technical replicates.

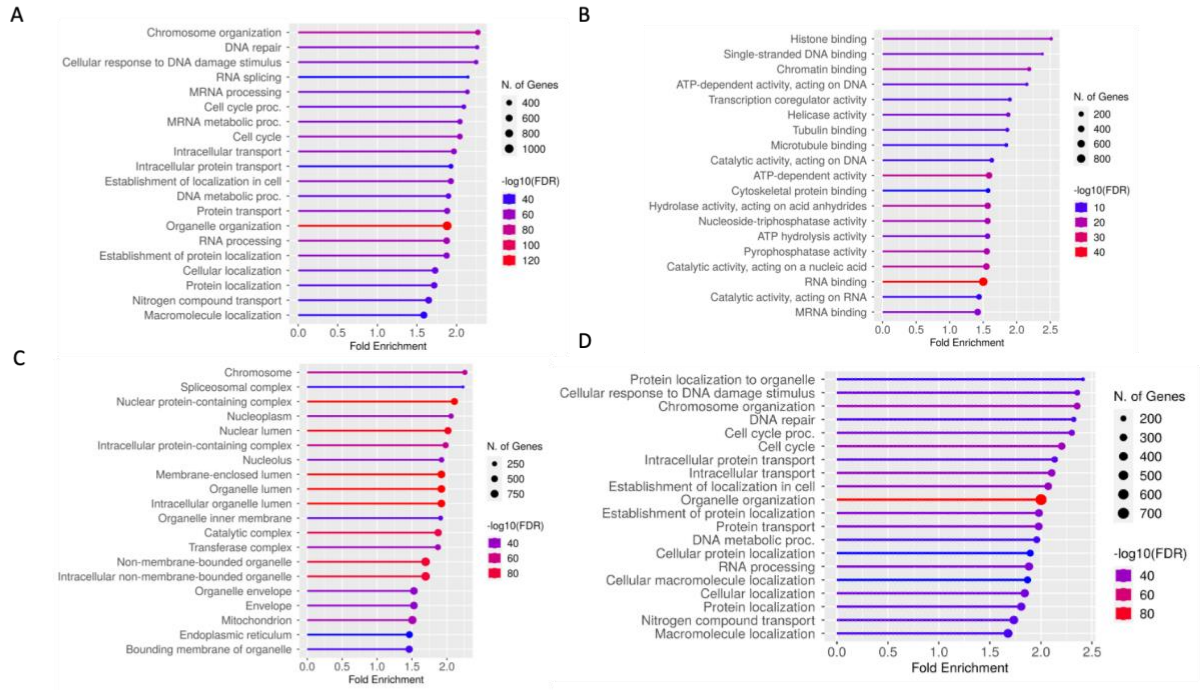


**Supplementary Figure 8. Functional categorisation of UBP5 target genes in ShinyGOv0.76 analysis.** A-C, GO analysis of all UBP5 target genes based on A, biological process; B, molecular function; and C, cellular component. D, GO analysis of UBP5 targets which gained H2Aub in *ubp5* based on biological process. False Discovery Rate (FDR) $< 0.05$.

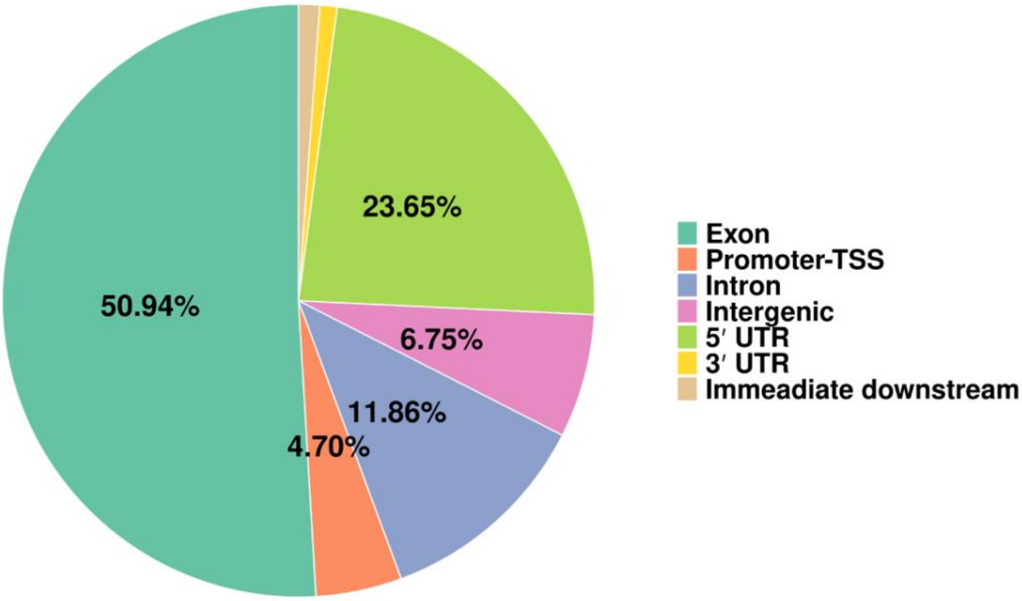


**Supplementary Figure 9. UBP5 majorly targets protein coding regions.** Pie chart showing the distribution of annotated genic and intergenic regions in the UBP5 binding peaks.

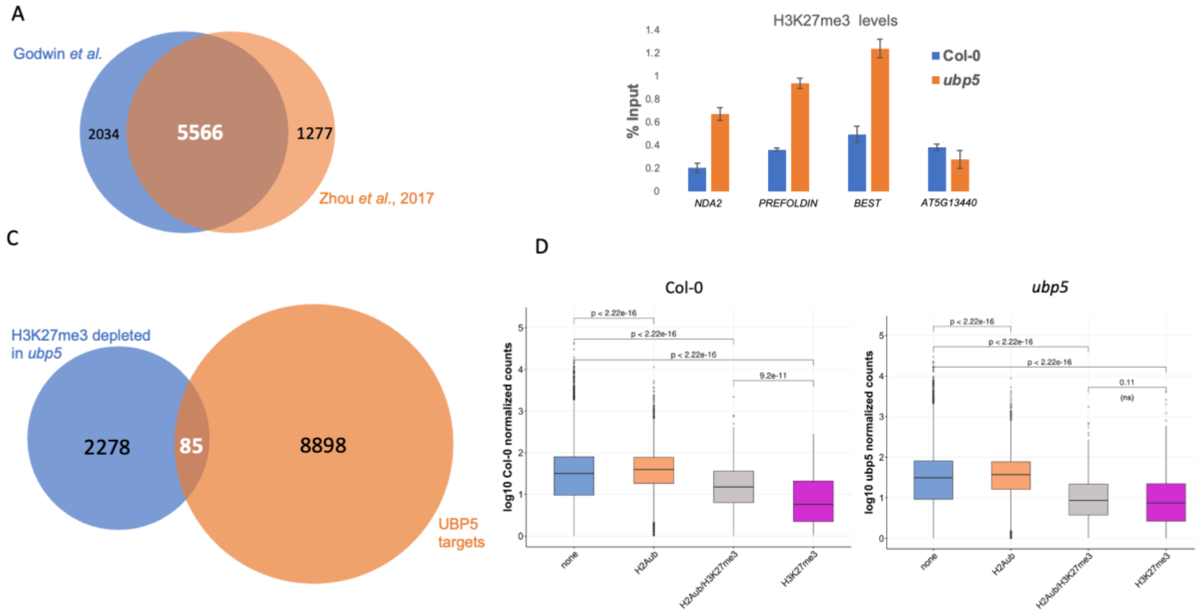


**Supplementary Figure 10. H2K27me3 changes in *ubp5*.** A, Venn diagram representing the overlap of H3K27me3 marked genes in Col-0 seedlings between our dataset and Zhou et al., 2017. B, ChIP-qPCR validation of H3K27me3 gained genes in *ubp5*. Selected loci *ALTERNATIVE NAD(P)H DEHYDROGENASE 2* (*NDA2* - AT2G29990), *PREFOLDIN* and *BEST* were among UBP5 targets transcriptionally downregulated in *ubp5*. *AT5G13440* was selected as negative control. ChIP-qPCR results shown as % of input DNA. Error bars indicate standard deviation of three technical replicates. C, Venn diagram representing H3K27me3 depleted genes in *ubp5* and UBP5 targets. D, Box plots showing expression levels of only-H2Aub, both H2Aub/H3K27me3 and only-H3K27me3 marked genes of Col-0 (left graph) and *ubp5* (right graph). "None" represents those genes lacking H2Aub and H3K27me3 marks. The statistical significance of the differences was calculated using the non-parametric one-sided Mann-Whitney-Wilcoxon test, p-values are indicated, ns represents non-significant. Results were generated using RNA-seq data from four biological replicates.

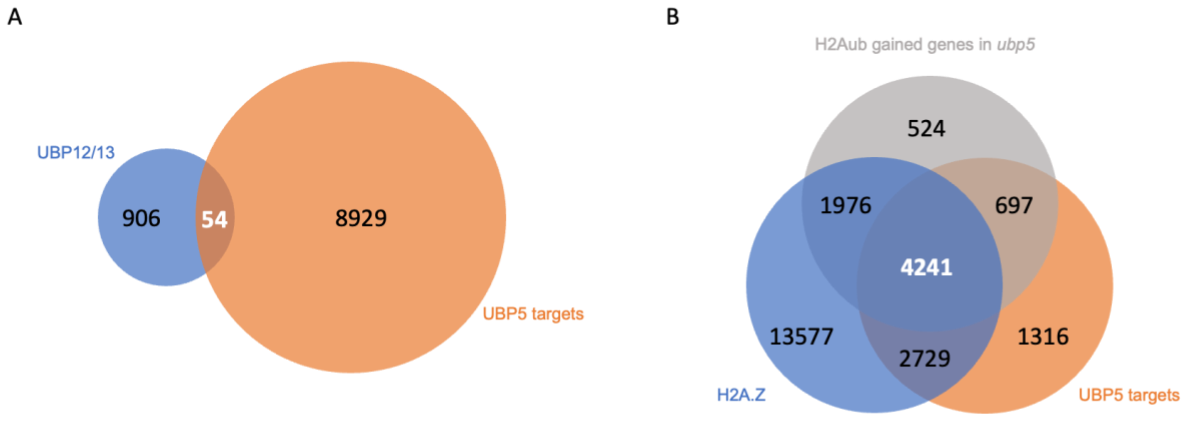


**Supplementary Figure 11. UBP5 shows low overlap with UBP12/13 and high overlap with H2A.Z.** A, Venn diagram showing the overlap between UBP12/13 regulated genes and UBP5 target genes. UBP12/13 regulated genes are defined as protein-coding genes gaining H2Aub and not losing H2Aub in the first 1kb of the gene body in *ubp12/13* mutants. UBP12/13 data from Kralemann *et al*., 2020. B, Venn diagram showing the overlap between UBP5 target genes, H2A.Z marked genes and H2Aub gained genes in *ubp5*. H2A.Z data from Wollmann *et al*., 2017.

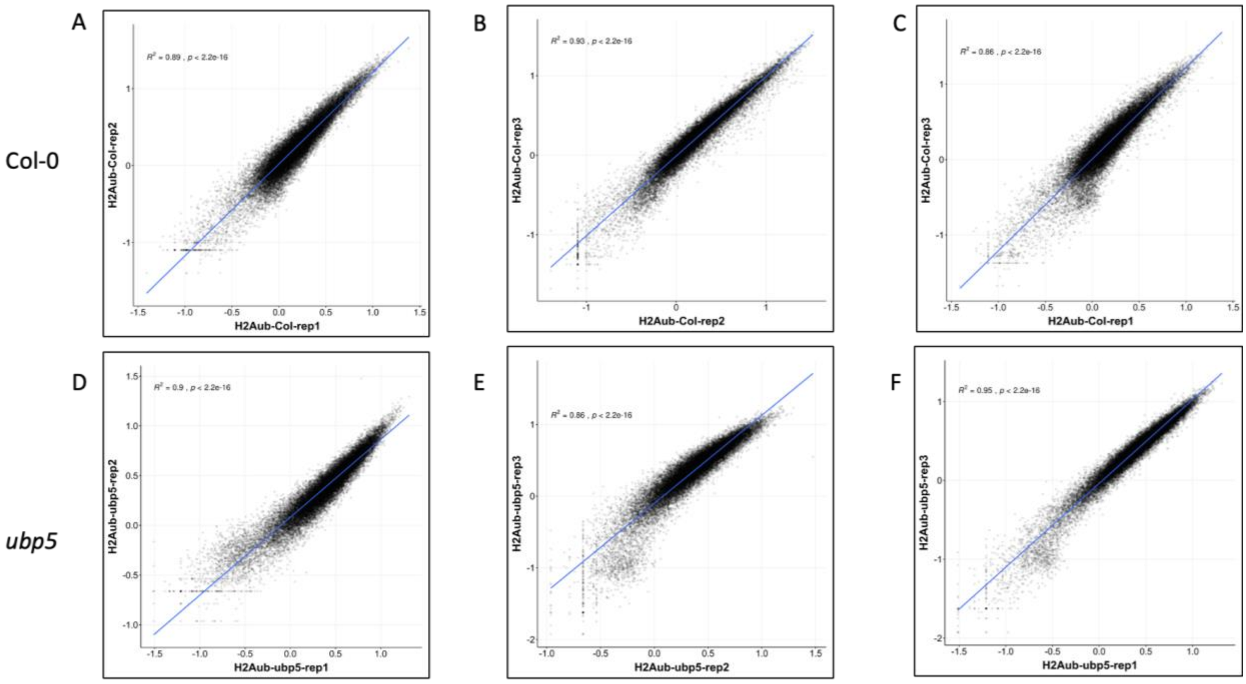


**Supplementary Figure 12. Comparison between three biological replicates for H2Aub ChIP-seq**. A-C, Comparison between Col-0 replicates. Each point represents the RPGC normalised counts across the first 1 kb of the gene body of protein coding genes. X and Y axes are in logarithmic scale (Log10). Lines are linear regression lines. D-F, Comparison between *ubp5* replicates. Coefficient of determination (R2) values and p values are mentioned on top of each panel.

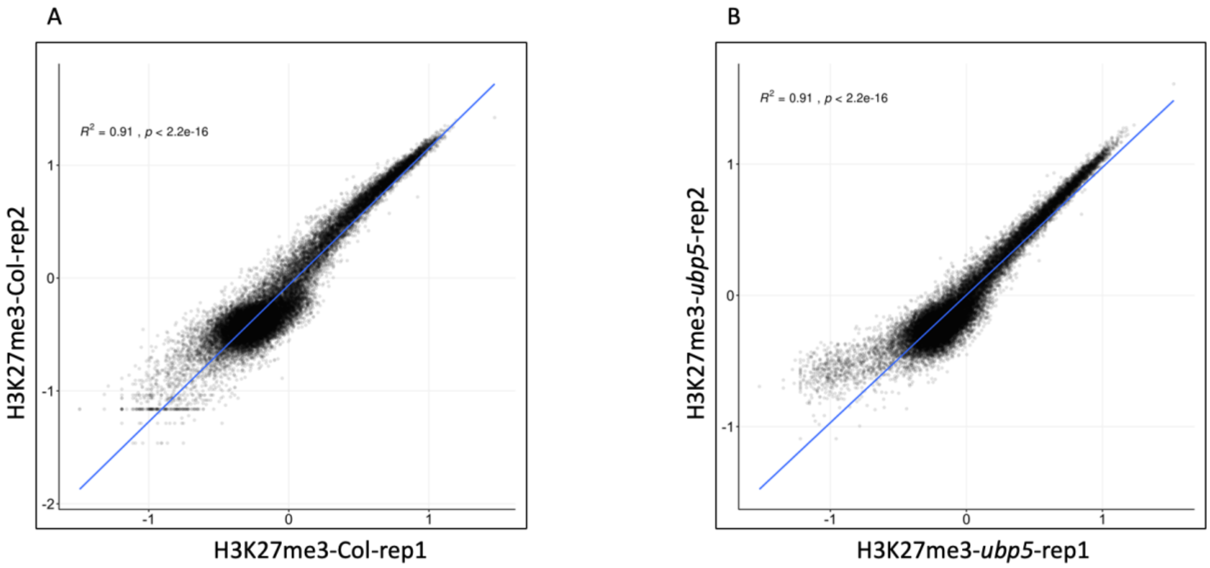


**Supplementary Figure 13. Comparison between two biological replicates for H3K27me3 ChIP-seq**. A, Comparison between Col-0 replicates. Each point represents the RPGC normalised counts across the first 1 kb of the gene body of a protein coding genes. X and Y axes are in logarithmic scale (Log10). B, Comparison between *ubp5* replicates. Lines are linear regression lines. Coefficient of determination (R2) values and p values were highlighted on top of each panel.

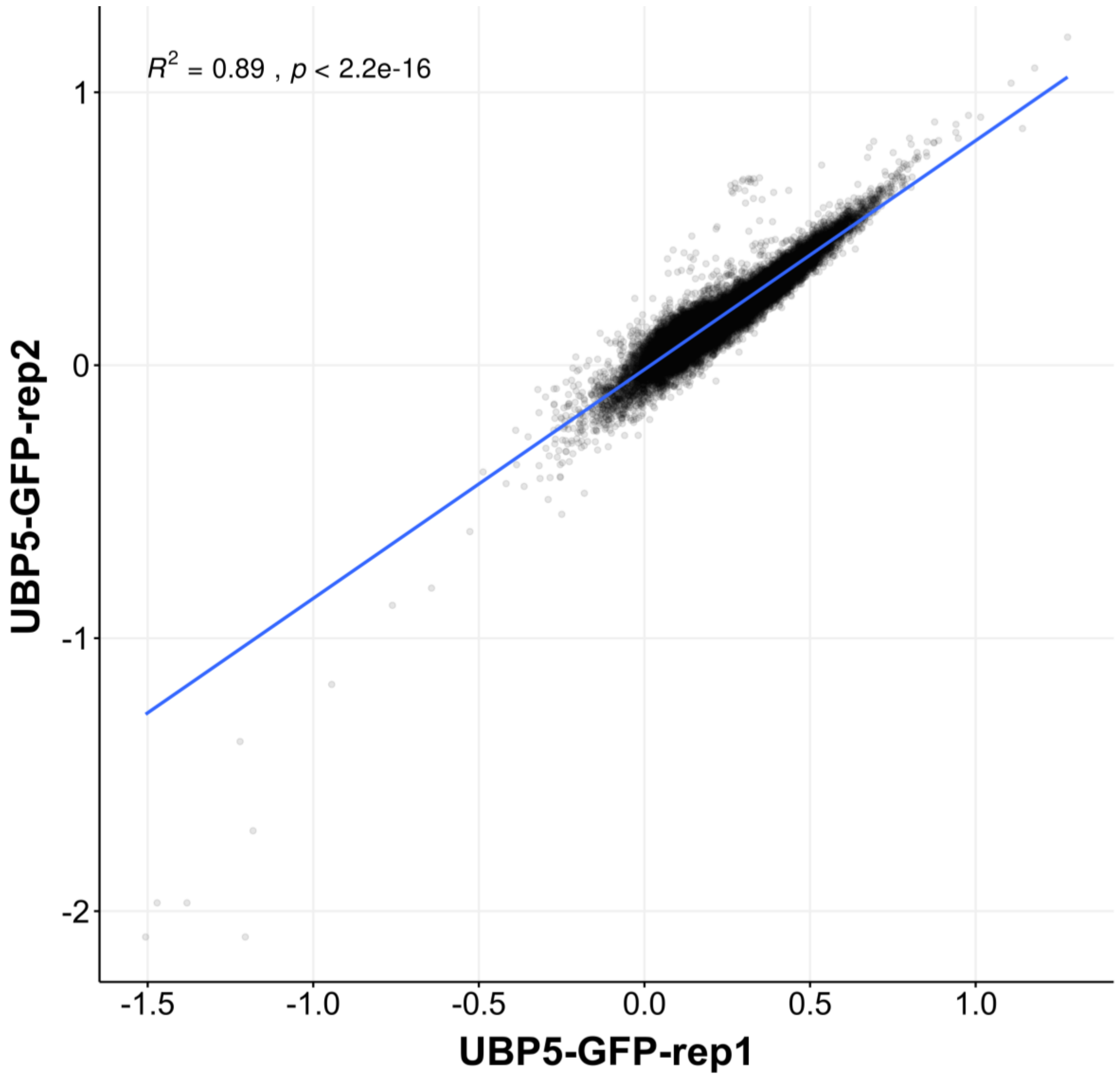


**Supplementary Figure 14. Comparison between two biological replicates of UBP5-GFP**. Each point represents the RPGC normalised counts across the first 1 kb of the gene body of a protein coding genes. X and Y axes are in logarithmic scale (Log10). Lines are linear regression lines. Coefficient of determination (R2) values and p values were mentioned on top of each panel.

**Supplementary table 1: Primers used in this study**

| **Primers designed to develop the CRISPR-Cas9 targeted mutagenesis specific vectors** | | |
|---|---|---|
| *UBP5_guide6-BsF* | ATATATGGTCTCGATTGTACGGGGGTGGTCCAACTCGTT | |
| *UBP5_guide6-F0* | TGTACGGGGGTGGTCCAACTCGTTTTAGAGCTAGAAATAGC | |
| *UBP5_guide20-R0* | ACCAGTCGGAGGAGTTGCTTTTCAATCTCTTAGTCGACTCTAC | |
| *UBP5_guide20-R0* | ATTATTGGTCTCGAAACAGTCGGAGGAGTTGCTTTTCAA | |
| **Primers for ChIP-qPCR** | | |
| Prefoldin subunit_Fw | AT1G03760 | ACCAACTACATTAAACGGGAGCA |
| Prefoldin subunit_Rv | AT1G03760 | ACAGTACGTCGACGAAAACGA |
| GAF1_Fw | AT5G59980 | CCACGAAAGCCATGGAGCTA |
| GAF1_RV | AT5G59980 | ATCGCGATGGAATCCGACAG |
| BEST_fw | AT3G61320 | CTCAAACCTCCCCGTGTCTC |
| BEST_Rv | AT3G61320 | TCGTCTGCCCAATCTGGAAC |
| H3K27me3 negative control | AT5G13440 | ACAAGCCAATTTTTGCTGAGC |
| H3K27me3 negative control | AT5G13440 | ACAACAGTCCGAGTGTCATGGT |
| H2Aub negative_Fw | Intergenic | CCATCTGCTCCACCGGGTAT |
| H2Aub negative_rv | Intergenic | GGATCGTAGAAGGCAGACCA |
| NDA2_Fw | | TTCTCTCGTCGGTGGCAAAC |
| NDA2_Rv | | TGCACAGCTCAAGAGACTCAG |
| AT4G39560_Fw | | TCGTAAAGCTCCGGTGAAGC |
| AT4G39560_Rv | | GACGTCGCCGATACTCTCAC |
| AT1G13880_Fw | | GATCTGGGCTCGTAGGTTGG |
| AT1G13880_Rv | | TCACGGATTAAATCGCGAAATC |
| **Primers for qRT-PCR** | | |
| *KNAT1*_Fw | AT4G08150 | CACATCCTCAACAATCCTGATGGG |
| *KNAT1*_Rv | AT4G08150 | TGGTTCTTGAGTTCCCGATCTTCG |
| *KNAT2*_Fw | AT1G70510 | CGTTCGACGAGGCTACAACTTTC |
| *KNAT2*_Rv | AT1G70510 | ACCGCACCATCATCTGAAAGAG |
| *STM*_Fw | AT1G62360 | ACCTTCCTCTTTCTCCGGTTATGG |
| *STM*_Rv | AT1G62360 | GCGCAAGAGCTGTCCTTTAAGC |
| *WUS*_Fw | AT2G17950 | TCATCACGGTGTTCCCATGCAG |
| *WUS*_Rv | AT2G17950 | CCCGTTATTGAAGCTGGGATATGG |
| *CLV3*_Fw | AT2G27250 | TAAGGACTGTTCCTTCGGGACCTG |
| *CLV3*_Rv | AT2G27250 | TCTTGGCTGTCTTGGTGGGTTC |
| *UBC21*_Fw | AT5G25760 | CTGCGACTCAGGGAATCTTCTAA |
| *UBC21*_Rv | AT5G25760 | TTGTGCCATTGAATTGAACCC |
| *UBP5*_Fw | AT2G40930 | CCAATCTGCAACAAGGTTTCTGT |
| *UBP5*_Rv | AT2G40930 | CTGTGACTGTTATTGCTCGTGT |
